# Functional ultrasound speckle decorrelation-based velocimetry of the brain

**DOI:** 10.1101/686774

**Authors:** Jianbo Tang, Dmitry D. Postnov, Kivilcim Kilic, Sefik Evren Erdener, Blaire Lee, John T. Giblin, Thomas L. Szabo, David A. Boas

## Abstract

A high-speed, contrast free, quantitative ultrasound velocimetry (vUS) for blood flow velocity imaging throughout the rodent brain is developed based on the normalized first order temporal autocorrelation function of the ultrasound field signal. vUS is able to quantify blood flow velocity in both transverse and axial directions, and is validated with numerical simulation, phantom experiments, and *in vivo* measurements. The functional imaging ability of vUS is demonstrated by monitoring blood flow velocity changes during whisker stimulation in awake mice. Compared to existing power Doppler and color Doppler-based functional ultrasound imaging techniques, vUS shows quantitative accuracy in estimating both axial and transverse flow speeds and resistance to acoustic attenuation and high frequency noise.

## 1. Introduction

Functional quantitative *in vivo* imaging of the entire brain with high spatial and temporal resolution remains an open quest in biomedical imaging. Current available methods are limited either by shallow penetration of optical microscopies that only allow imaging of superficial cortical layers, or by low spatiotemporal resolution such as functional magnetic resonance imaging or positron emission tomography. Ultrasound-based blood flow imaging techniques hold the promise to fulfill the unmet needs^[1,2]^, particularly with the emerging implementation of ultrafast ultrasound plane wave emission^[3]^ which paves the way for ultrasound to be applied for functional cerebral hemodynamic imaging of the entire rodent brain with 10-100 *μm* resolution.

Since the introduction of ultrafast plane wave emission-based Power Doppler functional ultrasound imaging (PD-fUS)^[4]^, an increasing number of studies are exploiting the capabilities of PD-fUS for functional brain imaging studies^[5–7]^. However, the exact relationship between the PD-fUS signal and the underlying physiological parameters is quite complex as the PD-fUS signal is also affected by the acoustic attenuation, beam pattern, clutter rejection and flow speed, in addition to the blood volume fraction and hematocrit^[8,9]^. On the other hand, ultrasound Color Doppler (CD-fUS) is able to measure a specific physiological parameter of the axial blood flow velocity but suffers from unstable estimations of mean speed due to the presence of noise and from incorrect estimation if opposite flows exist within the measurement voxel^[2,4,10–12]^. The microbubble tracking-based ultrasound localization microscopy (ULM^[13]^) method is able to map the whole mouse brain vasculature (coronal plane) and quantify the in-plane blood flow velocity (vULM^[13,14]^) with ~10 *μm* resolution. However, it suffers from a fundamental limitation of low temporal resolution as it requires extended data acquisition periods (~150 seconds for 75,000 images^[13]^) to accumulate sufficient microbubble events to form a single vascular image and corresponding velocity map, limiting its potential for functional brain imaging studies.

Here, we report a novel ultrasound speckle decorrelation-based velocimetry (vUS) method for blood flow velocity image of the rodent brain that overcomes the aforementioned limitations. We derived vUS theory which shows that the ultrasound field signal decorrelation in small vessels is not only determined by flow speed but also the axial velocity gradient and a phase term due to axial movement. We further developed a comprehensive experimental implementation and data processing methodology to apply vUS for blood flow velocity imaging of the rodent brain with high spatiotemporal resolution and without the need for exogenous contrast. We validated vUS with numerical simulations, phantom experiments, and *in vivo* measurements, and demonstrated the functional imaging ability of vUS by quantifying blood flow velocity changes during whisker stimulation in awake mice. We further show its advantage over PD-fUS and CD-fUS in terms of quantitative accuracy in estimating axial and transverse flow speeds and its resistance to acoustic attenuation and high frequency noise through phantom and *in vivo* measurements.

## 2. Results

### 2.1 vUS theory

The time varying ultrasound signal detected from a measurement voxel at time *t* can be considered as the integration of all moving point scatters within the voxel, and the ultrasound pressure arising from a given voxel can thus be written as,

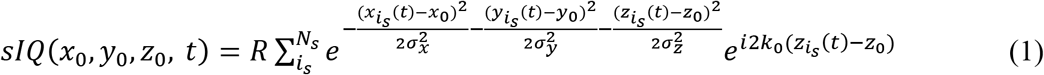

where, sIQ is the complex ultrasound quadrature signal of the moving particles of the voxel; *R* is the reflection factor; *i*_*s*_ is the index of the *i*^th^ scatterer; *N*_*s*_ is the total number of scatterers within the voxel; 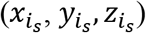 is the position of the *i*_*s*_ scatter; (*x*_0_, *y*_0_, *z*_0_) is the central position of the measurement voxel; *σ*_*x*_, *σ*_*y*_, and *σ*_*z*_ are the Gaussian profile width at the 1/*e* value of the maximum intensity of the point spread function (PSF) in *x, y*, and *z* directions, respectively; and *k*_0_ is the wave number of the central frequency of the transducer. In **Equation 1**, we assumed that all scatter points have the same reflection factor.

As shown in **Figure 1a**, the movement of particles will cause the detected ultrasound field signal to fluctuate in both magnitude and phase. This movement can be quantified based on the dynamic analysis theory of the normalized first-order field temporal autocorrelation function (*g*_1_(*τ*)). *g_1_(τ)* of a time varying ultrasound signal for a measurement voxel is given by,

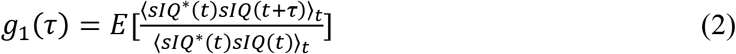

where, *τ* is the time lag; E[…] indicates the average over random initial positions; ⟨…⟩_*t*_ represents an ensemble temporal average; sIQ is the clutter rejected ultrasound quadrature signal; and * is the complex conjugate. **Figure 1b** illustrates the major characteristics of *g_1_(τ)*. Briefly, 1) *g_1_(τ)* decays faster for scattering particles flowing with higher speeds, 2) *g_1_(τ)* rotates and decays to (0, 0) in the complex plane, and 3) different flow angle has different decorrelation path in the complex plane, as shown in **Figure 1b2**. The rotating decorrelation in the complex plane is caused by the phase change due to axial movement. As shown in **Figure 1b3** flows with the same total speed but in different angles have the same magnitude decorrelation (left panel) but different ‘rotation paths’ in the complex plane (right panel). This feature gives *g_1_(τ)* analysis the ability to recover both axial velocity component and total flow speed.

**Figure 1.**
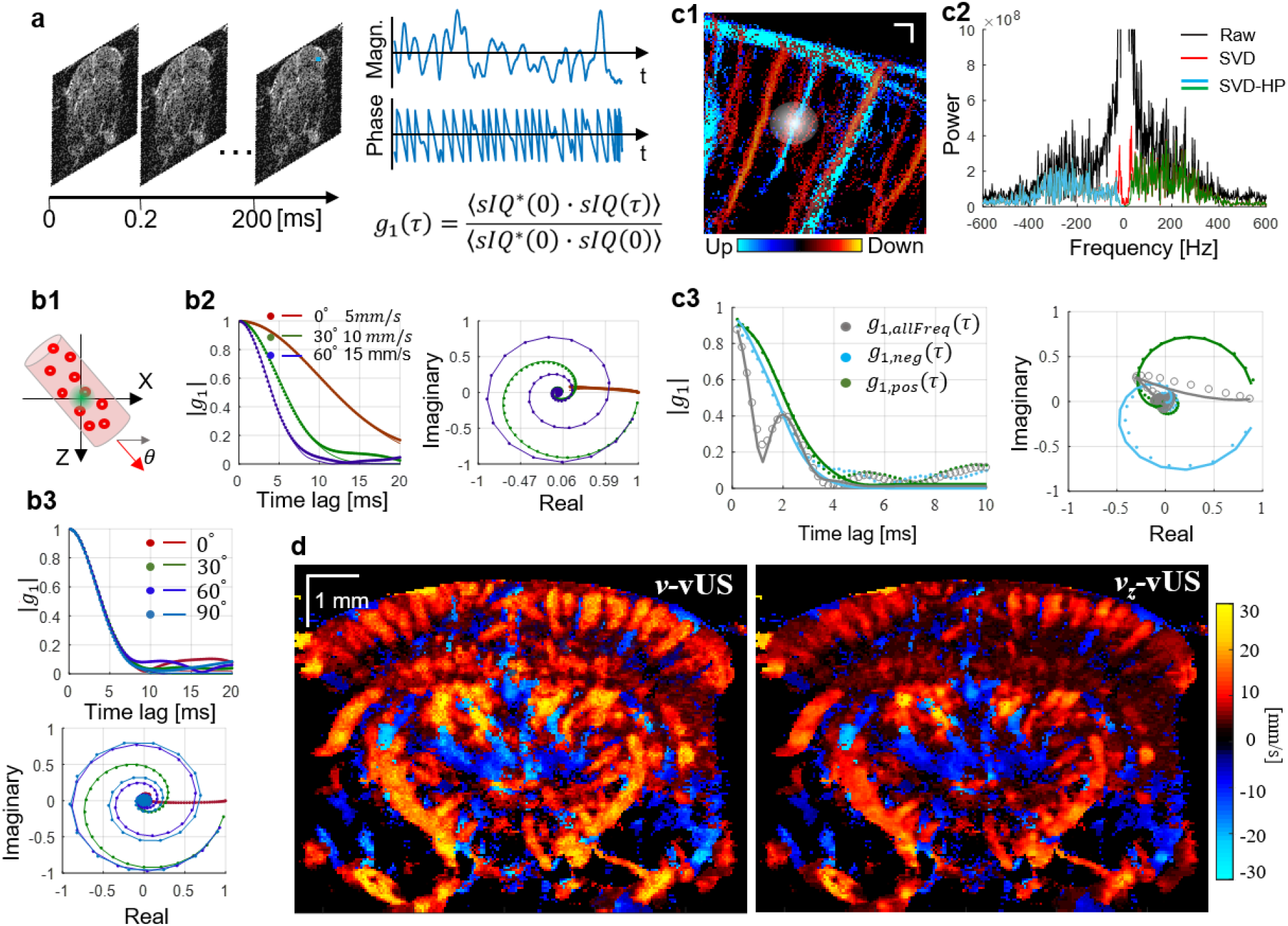
Principle of ultrasound field speckle decorrelation-based velocimetry (vUS). (a) A time series of a high frame rate complex ultrasound quadrature signal after bulk motion rejection *(sIQ(t))* was used for *g_1_(τ)* calculation. (b) Characteristics of *g_1_(τ)*; (b1) Scatterers flow through the measurement voxel at an angle θ; Magnitude decorrelation of |*g_1_(τ)*| and field decorrelation of *g_1_(τ)* in the complex plane at (b2) different angles with different speeds and (b3) different angles with the same speed (*v*_0_ = 15 mm/s). (c1) ULM measurement shows the microvasculature network in the brain; the white diffuse spot illustrates the ultrasound point spread function; (c2) Frequency power spectrum from *in vivo* data where descending and ascending vessels were observed in the same measurement voxel; (c3) *g_1_(τ)* calculated using whole frequency signal (gray circles), negative frequency signal (cyan dots), and positive frequency signal (green dots), respectively. (d) Representative total velocity map and axial velocity map reconstructed with vUS of a mouse brain; descending flow map is overlapped on the ascending flow map. The solid lines in (b&c) are the fitted *g_1_(τ)* using **Equation 3**.

When imaging the cerebral vasculature, the blood vessel diameter is usually less than the ultrasound system point spread function as indicated by **Figure 1c1**. In this case, the group velocity and velocity distribution must be taken into account as the relative movement of the scattering particles will result in additional decorrelation^[15]^. To simplify the derivation, we used a Gaussian speed distribution where, *v_gp_* is the group velocity; and *σ*_*v*_ describes the velocity distribution, and we finally arrive at,

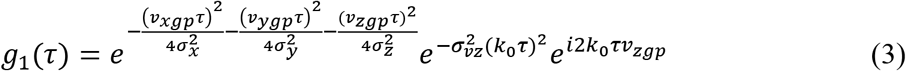

From **Equation 3**, we note that in addition to flow speed, the axial velocity distribution *σ*_*vz*_ also contributes to the magnitude decorrelation, and the axial velocity component leads to a phase term in *g_1_(τ)* decorrelation. For details regarding the theoretical derivation, please refer to the **Experimental Section-vUS theory derivation**.

In addition, we noticed from the *in vivo* data that it’s common to have opposite flows present in the same measurement voxel when imaging the rodent brain, as shown in **Figure 1c1**. In this case, *g_1_(τ)* is a mix of dynamics of opposite flows and behaves very differently from that of the single direction flow as can be observed from **Figs. 1b2** vs **c3** (gray circles). In addition, we observed that the majority of the mouse cerebral blood vessels contain an axial velocity component to the flow. This axial flow component causes the frequency spectrum to shift to negative values if the flow is away from the transducer, and positive if the flow is towards the transducer. Thus, we used a directional filter (positive-negative frequency separation) method to obtain the positive frequency and negative frequency signals for the *g_1_(τ)* calculation, as shown in **Figure 1c2**.

To implement the vUS technology, we developed a comprehensive vUS data acquisition and processing method (**Materials and Methods-vUS implementation** and **Figure S1**). **Figure 1d** shows representative in-plane total velocity and axial velocity maps of a mouse brain reconstructed by vUS. The descending flow velocity map which is reconstructed from the negative frequency component (*sIQ_neg_*) is overlapped on the ascending flow velocity map which is obtained from the positive frequency component (*sIQ_pos_*). Like the existing PD-fUS and CD-fUS techniques, vUS has an in-plane spatial resolution of ~100 *μm* which is determined by the ultrasound system acquisition parameters. **Figure S2** shows more vUS results at different coronal planes.

### 2.2. Validation of vUS

The numerical simulation validation (details in **Materials and Methods**) results shown in **Figure 2a** suggest that the vUS reconstructed total velocity (*v*), transverse velocity component (*v*_*x*_) and axial velocity component (*v*_*z*_) agree well with preset speeds and angles. It is worth noting that vUS is capable of measuring transverse flows (i.e. *θ* = 0°) and differentiating the axial velocity component from the transverse velocity component for the angled flows, as shown by results from flow angle *θ* = 30° and *θ* = 60°. For all simulation results, the correlation coefficient between *v*_*set*_ and *v_fit_mean_* were *r* >0.99 with p<0.001.

**Figure 2.**
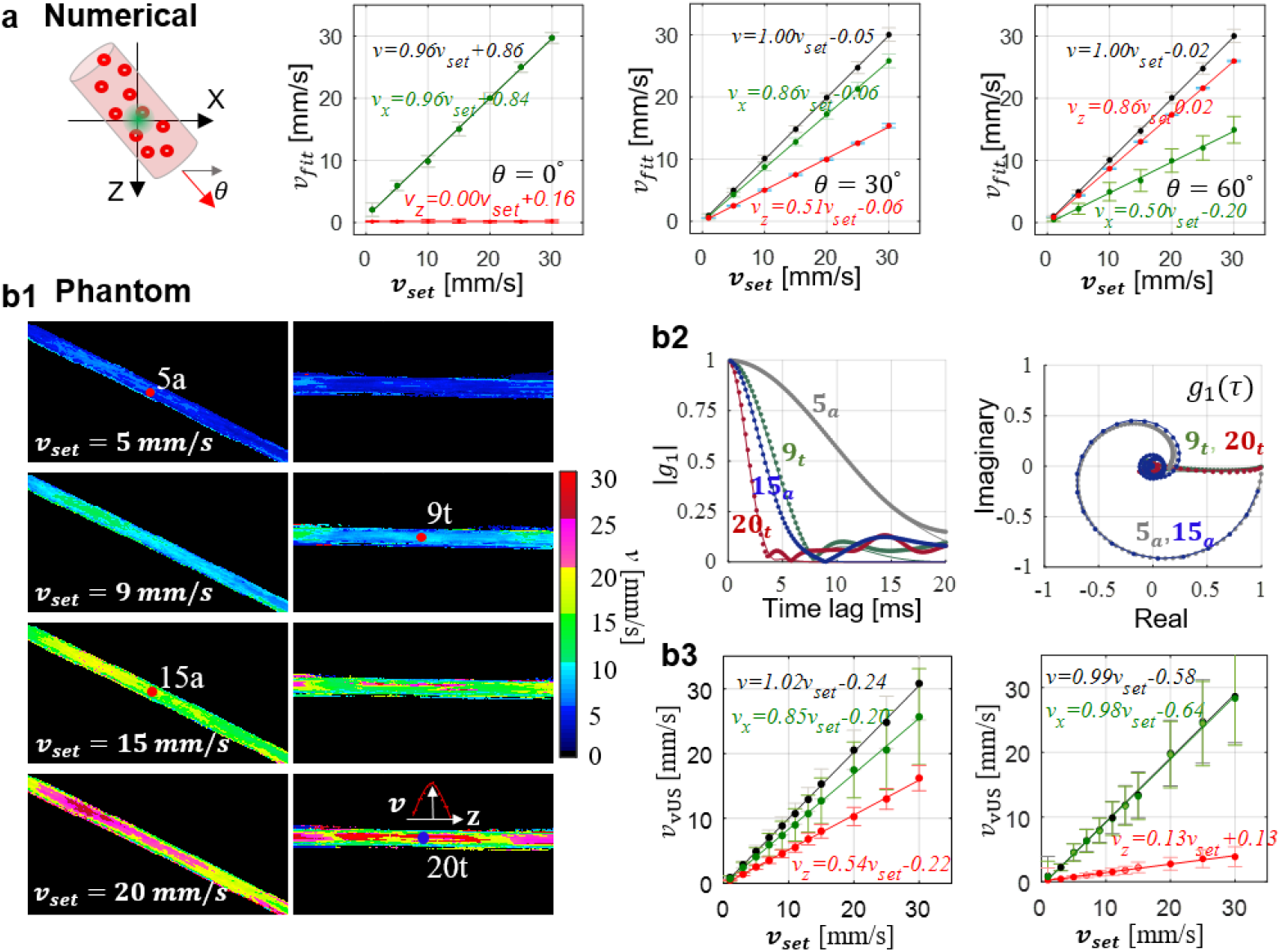
vUS numerical and phantom validation. (a) Numerical simulation validation with different flowing angles and speeds. Error bars: standard deviation. (b) Phantom validation of blood flowing through angled and transverse positioned micro tubes (inner diameter 580 *μm*). (b1) vUS reconstructed velocity maps of angled and transverse flows at different speeds. The inset in the right bottom panel shows the cross sectional laminar velocity profile of the transverse flow. (b2) Experimental *g_1_(τ)* (dots) and corresponding vUS fit results (solid lines) for both angled and transverse flows at different speeds. (b3) Results of vUS (*v*, *v*_*x*_ and *v*_*z*_) for transverse flow (*θ* ≈ 0°, left) and angled flow (*θ* ≈ 30°, right). Error bars: standard deviation.

The phantom validation experiments (details in **Materials and Methods**) were performed with blood samples flowing through a micro plastic tube buried within a static agarose phantom, as shown in **Figure 2b**. **Figure 2b1** shows the velocity maps of both angled and transverse flows at preset speeds of 5, 9, 15, and 20 mm/s. A laminar velocity profile was observed, particularly for higher flow speeds, as indicated in the inset of **Figure 2b1**. **Figure 2b2** shows the experimental (dots) and vUS fitted *g_1_(τ)*, from which we see that *g_1_(τ)* decays faster for higher speeds, and, as shown in the complex plane, *g_1_(τ)* rotates and decays to (0, 0) for angled flows (5_a_ and 15_a_) which is due to the axial velocity component inducing a phase shift as indicated in **Equation 3**. Different flow angles will have different ‘rotation paths’ in the complex plane. **Figure 2b3** shows the vUS reconstructed results compared to preset speeds, from which we note that the vUS measurements of total speed agree well with the preset speeds even for speeds as low as 1 mm/s for both transverse and angled flows. The correlation coefficient between *v*_*set*_ and *v*_*fit_mean*_ for transverse and angled flows were *r* >0.99 with p<0.001. **Figure S3** presents all phantom experiment results obtained with the vUS, CD-fUS, and PD-fUS analysis methods.

We further performed *in vivo* validation by comparing the velocity measured with ultrasound localization microscopy velocimetry (vULM, **Materials and Methods**) against vUS, as shown in **Figure 3.** We note that the measured axial velocity (**Figure 3a1**) and total velocity (**Figure 3b1**) agree well between vUS and vULM. The weighted scatter plots of all nonzero pixels between vUS and vULM in **Figure 3a2&b2** indicate that the vUS measurement is highly correlated with the vULM measurement. We further compared the mean velocity of 50 vessels marked in **Figure S4** between vULM and vUS. **Figure 3c1** shows the mean velocity and standard deviation measured with vULM (blue) and vUS (red) of the 50 vessels. **Figure 3c2** shows the scatter plot of the mean velocity of the 50 vessels measured with vULM and vUS. We note that the mean value of the 50 vessels agree well between vULM and vUS measurements with a linear relationship of 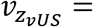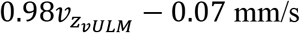, indicating the accuracy of vUS for *in vivo* blood flow velocity imaging within the rodent brain.

**Figure 3.**
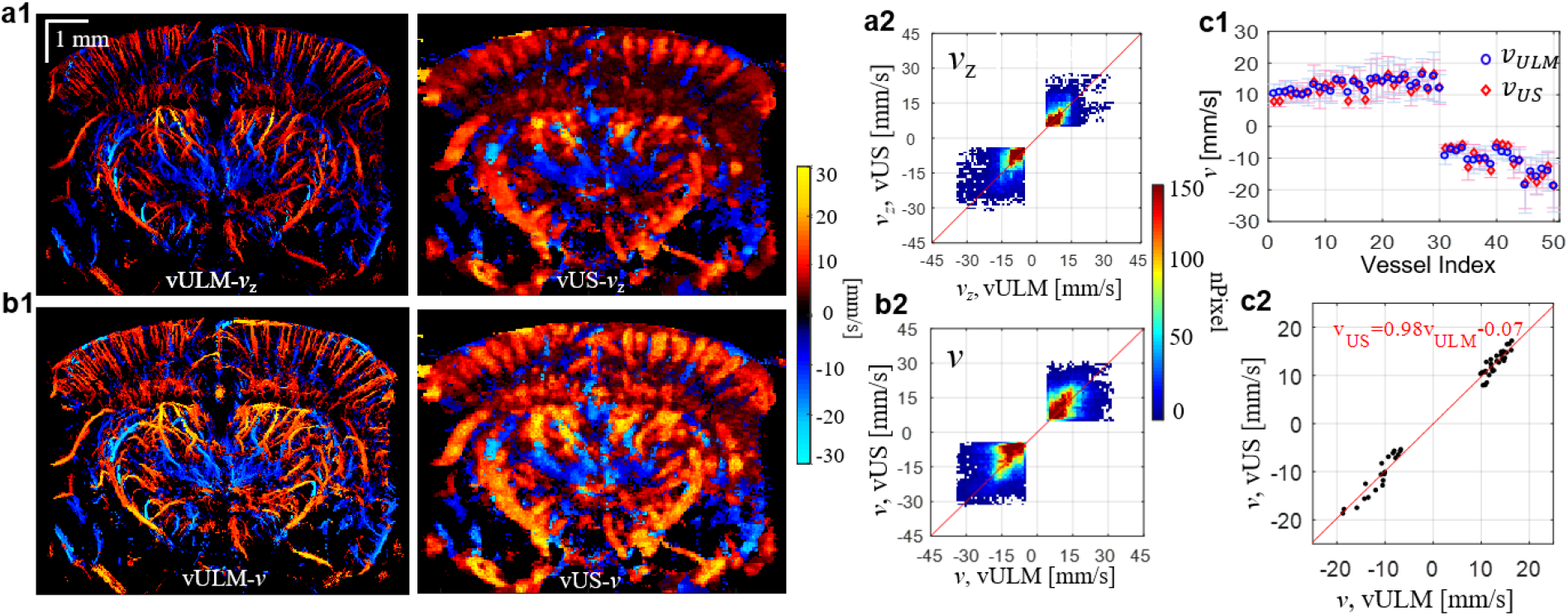
*in vivo* validation between vULM and vUS of axial velocity (a) and total velocity (b). (a2) and (b2) are pixel-to-pixel weighted scatter plot of common pixels of vULM and vUS with value |*v*| > 3 mm/s. (c1) Mean velocity and standard deviation measured with vULM (blue) and vUS (red) of 50 vessels marked in Supplementary Figure 4a. (c2) Cross correlation of the mean total velocity of the 50 vessels between vULM and vUS (r=0.984, p<0.001).

### 2.3. Blood flow velocity change evoked by whisker stimulation

To demonstrate the functional imaging capability of vUS, we measured the blood flow velocity response to whisker stimulation. We developed an animal preparation protocol using a polymethylpentene (PMP) film^[6]^ with a custom designed headbar for chronic ultrasound imaging in awake mice (**Materials and Methods**), as shown in **Figure 4a&b**. Following the published whisker stimulation protocol used in a previous PD-fUS study^[4]^, we used a stimulation pattern that consists of 30 s baseline followed by 10 trials of 15 s stimulation and with a 45 s interstimulus interval, as shown in **Figure 4c**. The vUS images were acquired at a rate of 1 frame/s.

**Figure 4.**
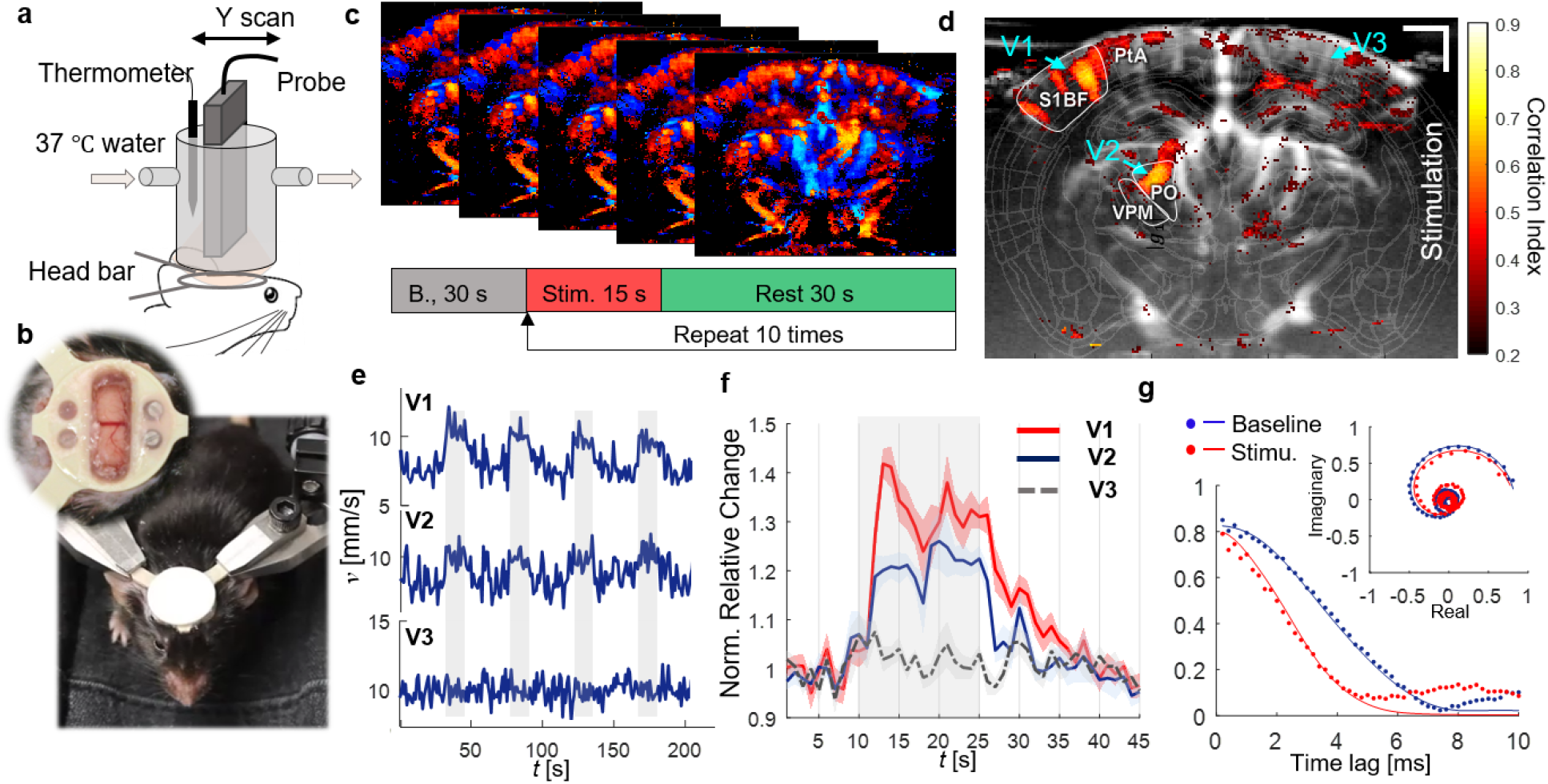
vUS of functional brain activation in awake mice. (a) Experimental setup. (b) Photos showing the trained mouse for awake-head fixed ultrasound imaging; inset: a PMP film protected cranial window was prepared in the center of the head bar for ultrasound imaging. (c) Whisker stimulation protocol and the vUS images were acquired at 1 frame/s. (d) Activation map in response to the mouse’s left whisker stimulation. S1BF: Primary somatosensory barrel field; PO: Posterior complex of the thalamus; VPM: Ventral posteromedial nucleus of the thalamus; PtA: Posterior parietal association. The ROIs were identified according to Allen Mouse Brain Atlas(*16*). (e) First 4 trials of blood flow velocity time course of vessels V1, V2, and V3 as marked in (d). The voxels of the three vessel ROIs were selected with absolute velocity value greater than 3 mm/s. Gray shades indicate when stimulation was on. (f) Average blood flow velocity relative change of the 10 trials for the three vessels. Error bar: standard error of the mean. (g) Representative *g_1_(τ)* from baseline (blue) and under stimulation (red) for the same pixel within V1. Solid lines: vUS fitted *g_1_(τ)*. Inset: *g_1_(τ)* in complex plane.

**Figure 4d** shows the correlation coefficient map between the blood flow velocity measured with vUS and the stimulation pattern. We note that in addition to the significant activation of vessels in the primary somatosensory barrel field (BF), the blood vessel flowing through the posterior complex (PO) and ventral posteromedial nucleus (VPM) of the thalamus also exhibited activation. Importantly, in addition to identifying significantly activated regions, vUS goes further and provides quantitative estimates of the evoked changes in the absolute flow velocity. The velocity time courses and velocity relative change averaged over the 10 trials of vessels V1 and V2 indicate robust blood flow velocity increases in response to the stimulation as shown in **Figs. 4e&f**. The time course of vessel V3 on the ipsilateral cortex of the stimulation was plotted as a control region, which shows no correlation with the stimulation. The **Supplemental Video 1** shows the relative blood flow velocity changes of the whole recording. We further compared the *g_1_(τ)* for baseline and under stimulation of the same spatial pixel in V1, as shown in **Figure 4g**. It is evident that *g_1_(τ)* decays faster when under stimulation compared to that during the baseline, indicative of faster dynamics, i.e. elevated blood flow speed in response to whisker stimulation. **Figure S5** shows more results of whisker stimulation experiments. Following the stimulation pattern commonly used in optical functional studies^[16]^, we used vUS to detect the cerebral blood flow velocity change in response to a 5 s whisker stimulation with a 25 s interstimulus interval, as shown in **Figure S5b**, and see that the measured blood flow velocity increases in response to the 5 s stimulation, indicating vUS is also sensitive to short duration stimulation evoked cerebral hemodynamic changes.

### 2.4. Comparison of vUS with PD-fUS and CD-fUS

The data set acquired for the vUS calculation can also be used for PD-fUS and CD-fUS data processing, so there can be a direct comparison of the different approaches. The advantages of vUS processing are apparent as shown in **Figure 5**. We see that 1) CD-fUS is only able to measure the axial velocity component (**Figure 5a**); 2) the signal intensity of PD-fUS is not linearly related to total speed but nonlinearly decreases with increasing speed (**Figure 5a2&b2**); and 3) vUS is able to measure the blood flow velocity of both angled (**Figure 5a**) and transverse (**Figure 5b**) flows and differentiate the axial velocity component from the transverse velocity component (**Figure 5a2**), indicating the advantages of vUS in quantitatively imaging flow speeds in both axial and transverse directions.

**Figure 5.**
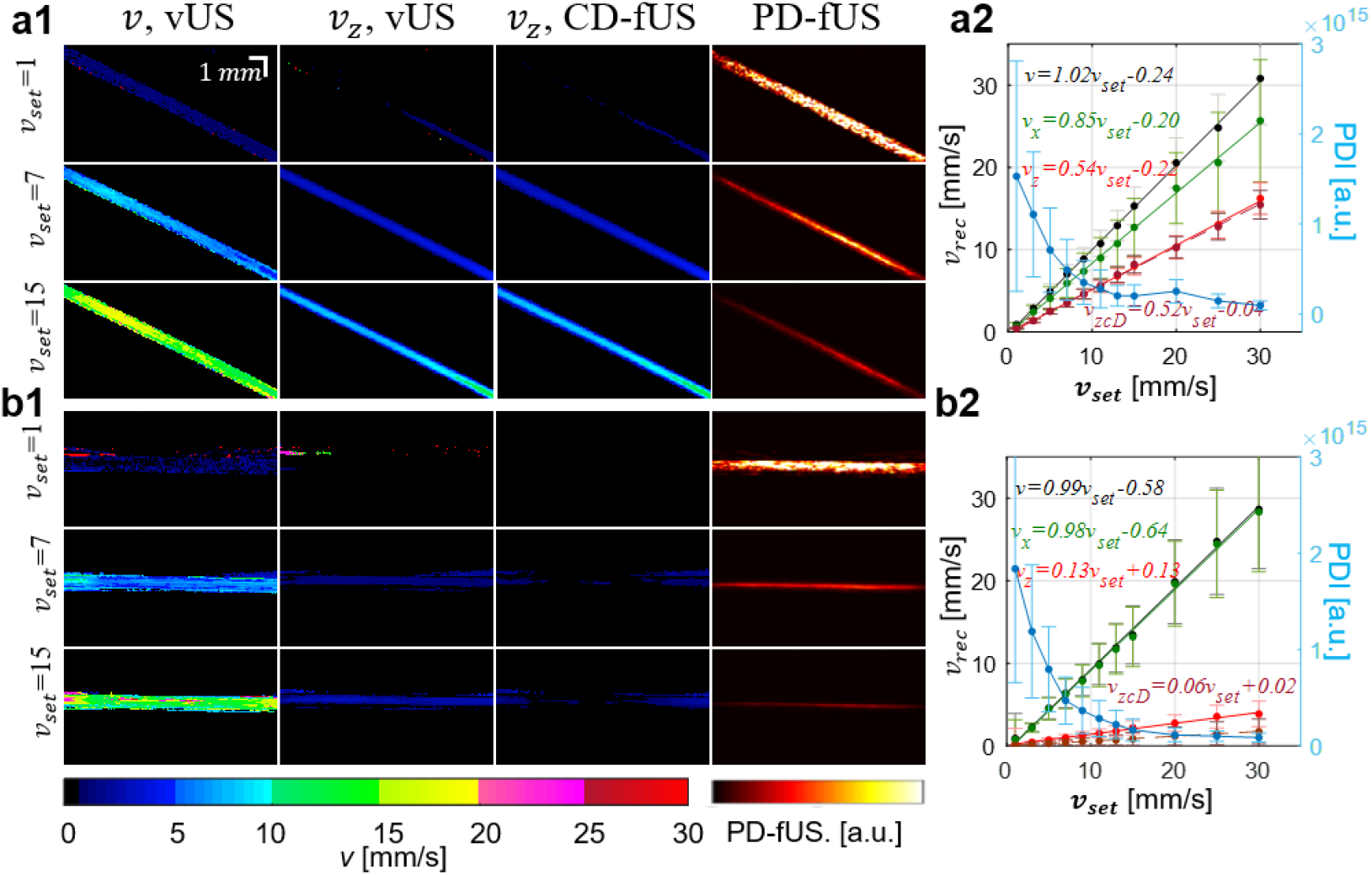
Phantom results comparison of vUS with Power Doppler-based fUS (PD-fUS) and Color Doppler-based fUS (CD-fUS). Angled (a) and transverse (B) flow phantom experiment results obtain with vUS (*v* and *v*_*z*_), CD-fUS (*v*_*z*_), and PD-fUS.

**Figure 6a** compares the *in vivo* measurements of ascending flow (positive frequency component) obtained with vUS and PD-fUS. Using the vULM measurement as the comparison standard of flow velocity, we note that vUS agrees well with vULM, while PD-fUS has high signal intensity in superficial layers and low signal intensity in deep regions, as indicated by the white and red arrows, indicating the strong dependence of the PD-fUS signal on acoustic attenuation. In contrast, vUS is not affected by acoustic attenuation as the normalization processing cancels the heterogeneous acoustic distribution. **Figure 6b1** shows the axial velocity maps obtained with conventional CD-fUS^[4]^ (**Online Methods**). The conventional CD-fUS suffers from underestimation of Doppler frequency (*fD*) due to mutual frequency cancellation when opposite flows exist within a measurement voxel, as illustrated in **Figure 6b2**. For a fair comparison between vUS and the Doppler methods, we applied CD-fUS processing on the directional filtered data that we used for vUS processing. As shown in **Figure 6c**, we note that the blood flow speed is overestimated by the directional filtering-based CD-fUS. This overestimation happens because of high frequency noise causing overestimation of the Doppler frequency (*f_D_*) when a directional filter is applied and thus a higher speed bias, as shown in **Figure 6c2**. In comparison, vUS doesn’t suffer from the high frequency noise as the high frequency noise is un-correlated and only causes *g_1_(τ)* to drop to a lower value at the first time lag but it doesn’t affect the decorrelation rate of *g_1_(τ)* at longer time lags, which is determined by the correlated motion of flowing red blood cells, as shown in the bottom panel of **Figure 6d2**. Thus, by fitting the decorrelation of *g_1_(τ)* the blood flow velocity can be accurately reconstructed by vUS, as shown in **Figure 6d1**.

**Figure 6.**
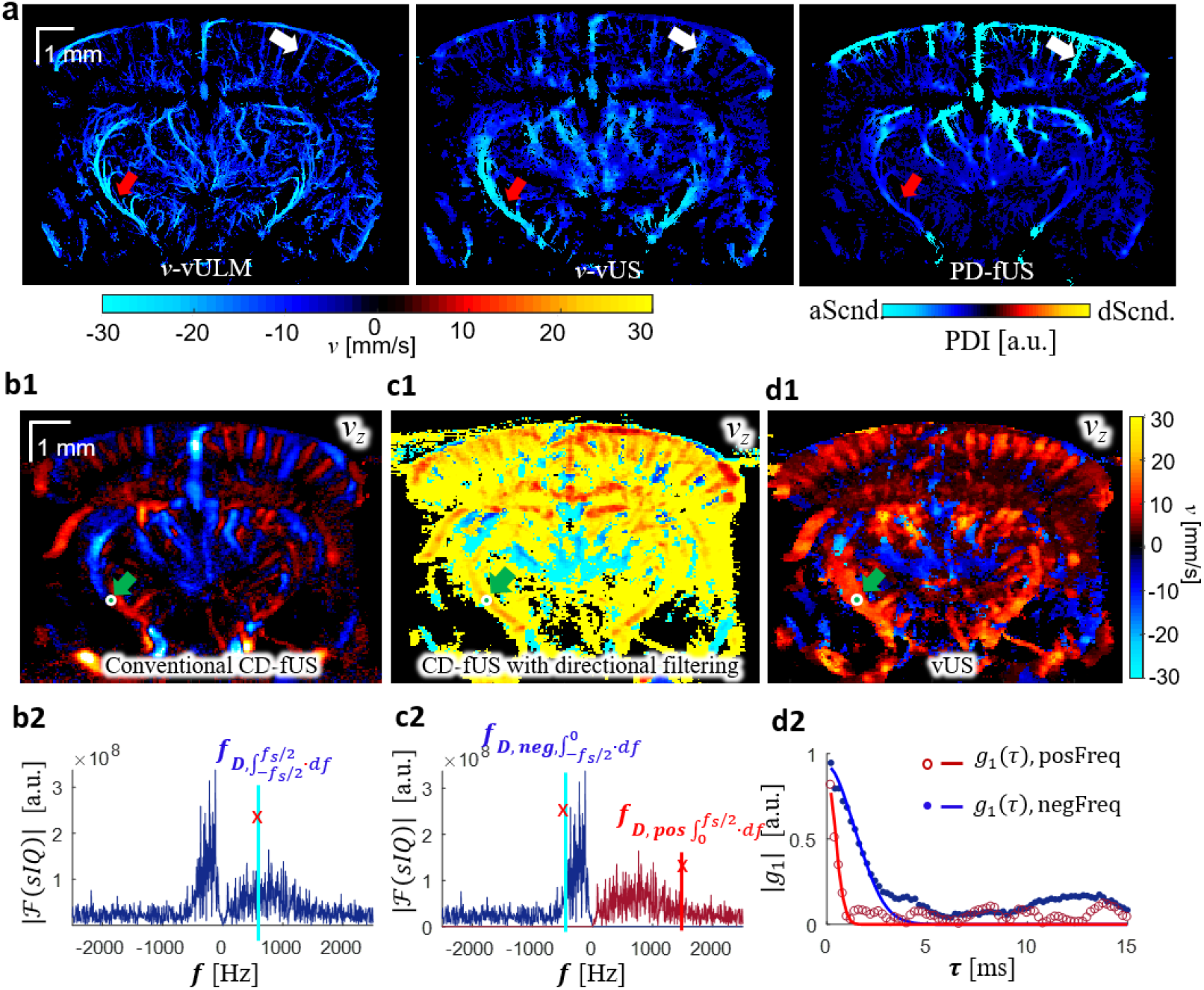
*in vivo* results comparison. (a) *in vivo* ascending flow results obtained with vULM, vUS, and PD-fUS, where vULM is used as the comparison standard and the ULM spatial mask was applied to both vUS and PD-fUS. (b1) Axial velocity (*v*_*z*_) map obtained with conventional CD-fUS; (b2) Doppler frequency (*f_D_*) is underestimated with conventional CD-fUS. (c1) Axial velocity map obtained with directional filtering-based CD-fUS; (c2) Doppler frequencies (*f_D,neg_* and *f_D,pos_*) are overestimated with the directional filtering-based CD-fUS. (d1) Axial velocity map obtained with vUS; (d2) *g_1_(τ)* calculated with positive frequency component and negative frequency component after directional filtering; dots: experimental data; solid line: theoretical fitting. Descending flow velocity maps were overlapped on ascending flow velocity maps in (c1) and (d1).

## 3. Discussion

The development of robust blood flow velocity measurement technologies has been of great importance in neuroscience research as quantifying blood flow alterations enables the assessment of brain disease^[17–19]^ and interpretation of regional neural function according to neurovascular coupling^[20]^. In this work, we introduced vUS based on the first-order temporal field autocorrelation function analysis of the ultrasound speckle fluctuations to quantify cerebral blood flow velocity with a temporal resolution of 1 frame/s (up to 5 frames/s in theory), with a greater than 10 mm penetration depth, and ~ 100 *μm* spatial resolution. vUS provides much deeper penetration compared to optical velocimetry methods which are usually restricted to superficial layers of less than 1 mm depth^[21]^ while maintaining high spatial and temporal resolution compared to magnetic resonance imaging-based phase contrast velocity mapping^[22]^.

Using ultrasound signal decorrelation analysis to estimate flow speed dates back to the 1970s. Atkinson and Berry^[23]^ have shown that the motion of moving scatterers is encoded in the fluctuations of the ultrasound signal and Bamber et al.^[24]^ demonstrated that the ultrasound signal decorrelation could be used to image tissue motion and blood flow. Wear and Popp and others^[8,9,25–28]^ showed that the decorrelation of ultrasound signal decays following a Gaussian form. In this paper, we showed that the ultrasound signal field decorrelation is governed by three terms, including the flow speed, the gradient of the axial velocity, and an axial velocity-dependent phase term. This phase term gives vUS the ability to differentiate the axial velocity component from the transverse velocity component.

The high frame rate ultrafast ultrasound plane-wave emission and acquisition paves the way for vUS implementation, which permits the speckle decorrelation caused by the moving scattering particles to be resolved with sufficiently high temporal resolution required to capture the speckle decorrelation within the small measurement voxels. The combination of spatiotemporal singular value decomposition and high pass filtering plays an important role in rejecting bulk motion which enables the decorrelation of *g_1_(τ)* to represent the dynamics of the motion of red blood cells and to not be confounded by bulk motion. For blood flow velocity imaging of the brain, vUS reconstructs both descending and ascending flow velocities from the negative frequency component and positive frequency component by applying directional filtering, respectively. We further developed a comprehensive fitting algorithm to reconstruct axial and transverse blood flow velocities. The proposed vUS technique was validated with numerical simulation, phantom experiments, and in vivo blood flow velocities obtained with vULM. The functional whisker stimulation experiment result agrees with previous rodent functional studies that mechanoreceptive whisker information reaches the barrel cortex via the thalamic VPM nuclei^[29]^, and the PO is a paralemniscal pathway for whisker signal processing^[30]^. This experiment demonstrates that vUS is sensitive to quantify the cerebral blood flow velocity change in response to functional stimulation and can be applied for brain imaging in awake mice.

Compared to PD-fUS (Power Doppler), vUS is a quantitative imaging modality for assessing blood flow velocity while the PD-fUS signal decreased with increasing speed and is strongly affected by the acoustic attenuation. Compared to CD-fUS (Color Doppler), vUS is able to measure both axial and transverse flow velocities and is resistant to high frequency noise compared to the directional filtering-based CD-fUS which suffers from large or random values in regions with a low signal-to-noise ratio. Compared to vULM, vUS has lower spatial resolution but has much higher temporal resolution (up to 5 Hz of vUS compared to 2 mins/frame of vULM) and is applicable for awake functional studies in rodents requiring high temporal resolution. In addition, it measures the flow velocity of the intrinsic contrast of red blood cells while vULM measures the speed of microbubbles. One important application that will be enabled by the absolute blood flow velocity measured with vUS is that the metabolic rate of oxygen can be quantitatively estimated if vUS measurements are combined with quantitative oxygenation measurements using multispectral photoacoustic tomography^[31,32]^, providing a new high resolution biomarker for neuroscience research.

A limitation is that vUS is not sensitive to measuring blood flow velocity in small vessels with low flow speeds due to the use of the spatiotemporal filter which rejects slow dynamics from the signal. Also, limited by the spatial resolution of the ultrasound system, the reconstructed blood flow velocity of a measurement voxel may represent integrated dynamics of multiple vessels that flow through the measurement voxel. For the results presented in this work, vUS was simplified to estimate in-plane 2D velocities (i.e., *v*_*x*_ and *v*_*z*_), ignoring decorrelation rom flow in the y-direction (see **Materials and Methods** for justification). This simplification, however, results in a moderate overestimation of the transverse velocity (*v*_*x*_) as *v*_*x*_ tends to compensate for the decorrelation caused by *v*_*y*_. Nevertheless, we note that the measured total velocity is very close to that obtained with vULM as shown in **Figure 3**. In the future, with the development of fast 3D ultrasound imaging technology using a 2D transducer matrix, vUS can be easily adopted for 3D velocimetry of the whole rodent brain.

## 4. Experimental Section

### 4.1. vUS theory derivation

The complex ultrasound quadrature signal of particles moving at the same speed in a measurement voxel can be written as,

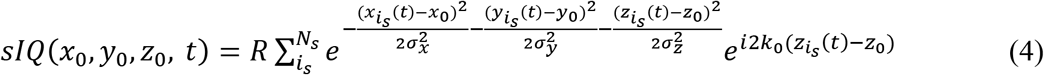

Considering the basic scenario that all scatters have identical dynamics, i.e. the scatters are moving in the same direction with same speed, the ultrasound pressure of the resolution voxel at time lag *τ* can be written as,

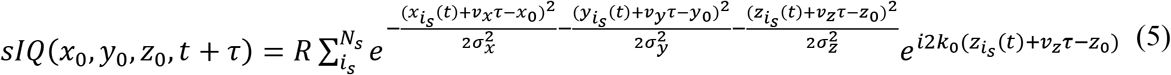

According to **Equation 2**, *g_1_(τ)* for particles flowing identically within the ultrasound measurement voxel can be derived to be,

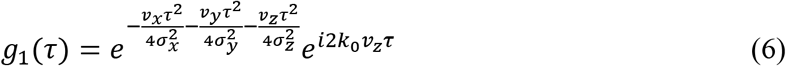

For microvasculature imaging of the rodent brain, the group velocity and velocity distribution must be taken into account as the relative movement of scatters will result in additional decorrelation. To simplify the derivation, we used a Gaussian distributed velocity model to describe the velocity distributed flow,

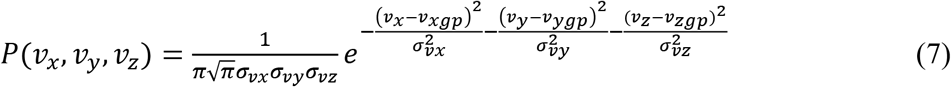

where, *P*(*v*_*x*_, *v*_*y*_, *v*_*z*_) is the velocity distribution probability; *v*_*gp*_ is the group velocity; and *σ*_*v*_ describes the velocity distribution.

*g_1_(τ)* for the Gaussian speed distribution flow is derived to be,

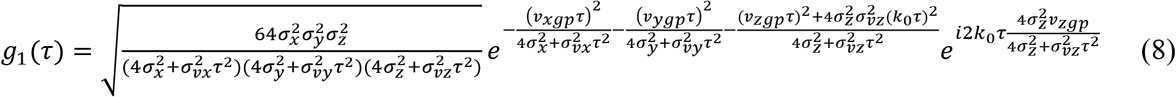

From our observations, the typical decorrelation time (*τ*_*c*_) for blood flow with a speed around 10 mm/s is ~5 ms. Therefore, 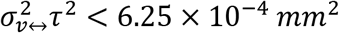 which is more than 8 times smaller than 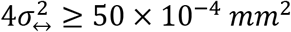, where ‘↔’ represents the coordinate direction (i.e., *x, y* or *z*). Thus, the theoretical equation of *g_1_(τ)* can be further simplified to be,

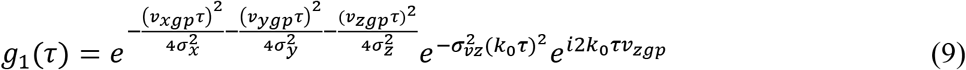

where, *σ*_*x*_, *σ*_*y*_, and *σ*_*z*_ are the Gaussian profile width at the 1/*e* value of the maximum intensity of the point spread function (PSF) in *x, y*, and *z* directions, respectively; *v*_*gp*_ is the group velocity; and *σ*_*vz*_ describes the axial velocity distribution; and *k*_0_ is the wave number of the central frequency of the transducer.

### 4.2. vUS implementation

#### 4.2.1. Coherent plane wave compounding-based data acquisition

The ultrasound signal was acquired with a commercial ultrafast ultrasound imaging system (Vantage 256, Verasonics Inc. Kirkland, WA, USA) and a linear ultrasonic probe (L22-14v, Verasonics Inc. Kirkland, WA, USA). The Vantage 256 system has 256 parallelized emission and receiving channels, and can acquire planar images at a frame rate up to 30 kHz when the imaging depth is ~15 mm. The L22-14v ultrasonic probe has 128 transducer elements with a pitch of 0.1 mm and a center frequency of 18.5 MHz with a bandwidth of 12.4 MHz (67%, −6 dB). It has an elevation focus at z=6 mm.

To ensure sufficient temporal resolution, the ultrasound plane wave frame rate was set to 30 kHz which was mainly limited by the transmit time of the ultrasound signal in the sample through the intended imaging depth, as shown in **Figure S1a**. To enhance the signal-to-noise ratio while preserving sufficient temporal resolution, we further employed coherence plane wave compounding^[33]^ at five emitting angles (−6°, −3°, 0°, 3°, 6°) to form a compounded image whose frame rate was 5 kHz, as shown in **Figure S1b**.

In addition, to acquire sufficient ensemble averaging of the US speckle fluctuations for the vUS analysis, we acquired 200 ms of data, i.e. 1,000 compounded images, to calculate *g_1_(τ)* over a range of 0<τ<20 ms. Therefore, the maximum vUS frame rate is 5 frames/s. However, for extended data acquisition (i.e. >1 mins) the maximum vUS frame rate was reduced to 1 frame/s due to limit data transfer and saving requirements.

#### 4.2.2. Clutter rejection

For the phantom data processing, we used a spatiotemporal filtering method (singular value decomposition, SVD, **Equation 10**^[34]^) to remove the first two (*Nc*=3) highest singular value signal components. To reject the bulk motion signal from the *in vivo* data, we used a combination of SVD and high pass filtering. The first 20 highest singular value signal components were removed (*Nc*=21), followed by a fourth order Butterworth high pass filtering with a cutoff frequency of 25 Hz corresponding with a 1 mm/s speed cutoff.

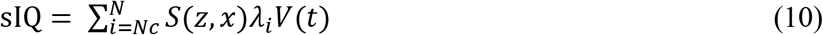

where, sIQ is the dynamic signal; *Nc* is the cutoff rank for SVD processing; *S*(*z*, *x*) is the spatial singular matrix; *λ*_*i*_ is the singular value corresponding with the *i*^th^ rank; and *V*(t) is the temporal singular vector.

#### 4.2.3. vUS fitting algorithm

**Figure S1d** summarizes the vUS data processing algorithm. Based on the developed vUS theory for *in vivo* brain imaging, the clutter rejected sIQ data of a measurement voxel, sIQ(z, x), was first directionally filtered to obtain the negative frequency signal component (descending flow) and the positive frequency signal component (ascending flow) using the directional filtering processing (**Equation 11&12**).

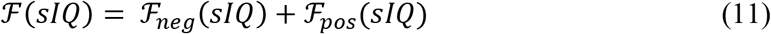

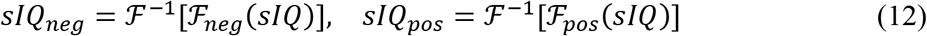

where, *sIQ*_*neg*_ and *sIQ*_*pos*_ are the complex ultrasound quadrature signal of the negative frequency and positive frequency, respectively; ℱ denotes the Fourier transform; and ℱ^−1^denotes the inverse Fourier transform. 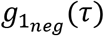 and 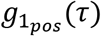 for *sIQ*_*neg*_ and *sIQ*_*pos*_ are obtained using **Equation 2**, respectively.

We used criteria including the ratio of positive/negative frequency power to whole frequency power (**Equation 13**) and the absolute value of *g_1_(τ)* at the first time lag (**Equation 14**) to control signal quality for data processing.

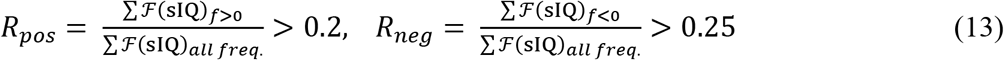

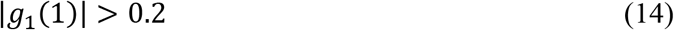

where, ℱ denotes the Fourier transform. These criteria enable us to skip the poor quality data, which also greatly reduces the processing time.

Then, the fitting procedure is applied for both *sIQ*_*neg*_ and *sIQ*_*pos*_, respectively. In practice, random noise results in a prompt ‘drop’ of *g*_1_(1), i.e. the change of *g*_1_(0) to *g*_1_(1) is not a smooth transition compared to *g*_1_(1) to the end of the decorrelation as the noise is uncorrelated. We therefore modified the *g_1_(τ)* equation by using an ‘F’ factor to account for this ‘drop’. Also, it is worth noting that when using a linear transducer array the ultrasound PSF is anisotropic in the transverse directions, i.e. *σ*_*x*_ ≠ *σ*_*y*_. In our experimental setup, *σ*_*y*_ was more than 3 times larger than *σ*_*x*_ which results in a more than 9 times slower signal decorrelation rate from *v*_*ygp*_ compared to that from *v*_*xgp*_. Therefore, we omitted the *y* component from the *g_1_(τ)* fitting to simplify the data processing. In addition, in the case of Gaussian velocity distribution, *σ*_*vz*_ is proportional to the maximum speed in the center line and also linearly related to the group velocity *v*_*zgp*_. Thus *σ*_*vz*_ in **Equation 3** can be replaced with *σ*_*vz*_ = *p* ∙ *v*_*zgp*_ where *p* is a linear factor with a range of [0 1]. Thus the theoretical *g_1_(τ)* model used for fitting the experimental data is,

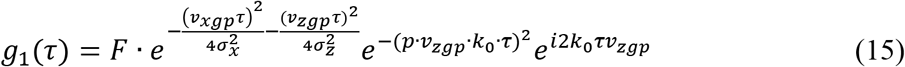

where *F* represents the correlated dynamic fraction which accounts for the *g_1_(τ)* value drop at the first time lag due to uncorrelated signal fluctuations (e.g. noise); *v*_*x*_ and *v*_*z*_ are the flow speed in the *x* and *z* directions respectively; *σ*_*vz*_ = *p* ∙ *v*_*z*_ accounts for the speed distribution within the measurement voxel where *p* is a linear factor with a range of [0 1]; *σ*_*x*_ and *σ*_*z*_ are the US voxel Gaussian profile width at the 1/*e* value of the maximum intensity of the point spread function (PSF) in the *x* and *z* directions, respectively; and *k*_0_ = 2π⁄*λ*_0_ is the wave number of the central frequency of the transducer.

A proper initial guess of the unknown parameters (i.e., *F*, *v*_*xgp*_, *v*_*zgp*_, and *p*) is important to achieve high fitting accuracy and efficiency. The initial guess of *F*_0_ was set to be *F*_0_=|*g*_1_(1)|. As the axial movement caused the phase change of *g_1_(τ)*, we used the phase information of *g_1_(τ)* to determine *v*_*zgp0*_ by finding the time lag *τ_v_* when *g_1_(τ)* reaches the first minimum.

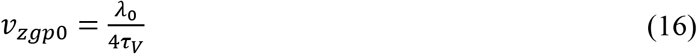

We tested a mesh of *v*_*xgp*_ and *p* values to determine the initial guess of *v*_*xgp0*_ and *p*_0_ by finding the pair of *v*_*xgp0*_ and *p*_0_ that maximizes the coefficient of determination, R. R is defined in **Equation 17** and was also used in the final fitting process as the objective function for a constrained least squares regression non-linear fitting procedure to estimate the values for *F*, *v*_*xgp*_, *v*_*zgp*_, and *p* based on the initial guesses.

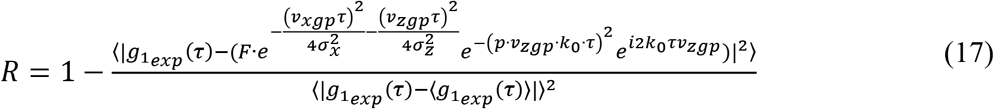

where, 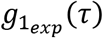 is the experimental *g_1_(τ)* calculated with **Equation 2**; 〈… 〉 indicates temporal ensemble averaging; and |…| indicates the absolute value.

Finally, the axial and total velocity maps were obtained for both descending and ascending flows, as shown in **Figure S1e**.

### 4.3. Power Doppler-fUS and Color Doppler-fUS calculation

The Power Doppler image (PD-fUS) was calculated as^[4]^,

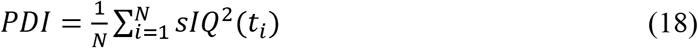

where, *N* is the number of samples and sIQ is the complex ultrasound quadrature signal of the moving particles.

The axial velocity based on the conventional Color Doppler calculation is obtained with^[10]^,

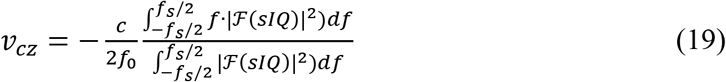

where, *c* is the sound speed in the medium and *c*= 1540 m/s was used in this study; *f*_0_ is the transducer center frequency; *f*_*s*_ is the frame rate; and ℱ denotes the Fourier transform.

Further, for a fair comparison with vUS which obtains velocity map based on the directional filtered data (*sIQ*_*neg*_ and *sIQ*_*pos*_), we used Color Doppler to process the same directional filtered data to obtain descending and ascending speeds (**Figure 6c1**),

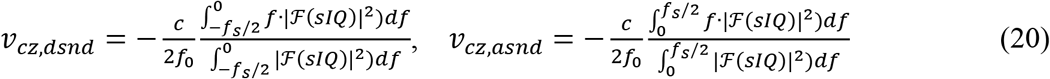

### 4.4. Ultrasound Localization Microscopy

The ultrasound localization microscopy (ULM) images and the ULM-based velocity maps (vULM) were obtained based on a microbubble tracking and accumulation method described in^[13,14]^. Briefly, a frame-to-frame subtraction was applied to the IQ data to get the dynamic microbubble signal. The images of the microbubble were rescaled to have a pixel size of 10 *μm* × 10 *μm*. The centroid position for each microbubble was then identified with 10 *μm* precision by deconvolving the system point spread function. By accumulating the centroid positions over time, a high resolution image of the cerebral vasculature image (ULM) is obtained. Further, by identifying and tracking the same microbubble’s position, the in-plane flow velocity of the microbubble can be calculated based on the travel distance and the imaging frame rate. The final velocity for coordinates (*z,x*) consists of descending and ascending flows, and the speed for each direction was obtained by averaging the same directional flow speed at all time points when the absolute value was greater than 0, respectively.

### 4.5. Numerical Simulation

In this study, two dimensional (x-z) flow and ultrasound detection was simulated to validate vUS. Point scattering particles (5 *μm* in diameter) were randomly generated at the initialization segment which is outside the ultrasound measurement voxel. Then the flowing positions were calculated for all time points based on the preset flow speed and flow angle at a temporal rate of 5 KHz. The detected ultrasound signal (*sIQ*) was obtained based on **Equation 1** for each time point. Then the simulated *g_1_(τ)* was calculated according to **Equation 2** with 1000 observation time points (i.e. 200 ms) and 100 autocorrelation calculation time lags (i.e. 20 ms). Flow velocity was then reconstructed by applying vUS processing on the simulated *g_1_(τ)*.

### 4.6. Phantom experiment and data processing

For the phantom validation experiment, a plastic micro tube (inner diameter 580 *μm*, Intramedic Inc.) was buried in a homemade agar phantom with an angle of ~ 30° (angled flow), and another plastic micro tube was aligned close to ~ 0° (transvers flow) in another homemade agar phantom. A blood solution was pumped through the tubes with a syringe pump (Harvard Apparatus) at speeds of 1, 3, 5, 7, 9, 11, 13, 15, 20, 25, and 30 mm/s. SVD was performed to filter the background signal clutter by removing the first two highest singular value components. Since the diameter of the tube is much larger than the ultrasound resolution, the red blood cell speed distribution can be considered uniform. Therefore, the linear value *p* in **Equation 15** was set to 0 (i.e. *σ*_*vz*_ = 0) for the phantom data processing.

### 4.7. Animal preparation

The animal experiments were conducted following the Guide for the Care and Use of Laboratory Animals, and the experiment protocol was approved by the Institutional Animal Care and Use Committees of Boston University.

In this study, 12-16-week old C57BL/6 mice (22-28g, male, Charles River Laboratories) were used. Animals were housed under diurnal lighting conditions with free access to food and water. Mice were anesthetized with isoflurane (3% induction, 1–1.5% maintenance, in 1L/min oxygen) while the body temperature was maintained with a homeothermic blanket control unit (Kent Scientific) during surgery and anesthetized imaging sessions. After removal of the scalp, a custom-made PEEK headbar was attached to the skull using dental acrylic and bone screws. The skull between lambda and bregma extending to temporal ridges was removed as a strip. A PMP film cut to the size of the craniotomy was then secured to the skull edges. Since the PMP is flexible, brain is protected by a cap attached to the head bar. The animal was allowed to recover for 3 weeks before the imaging sessions. During surgery and anesthetized imaging, heart rate and oxygen saturation was non-invasively monitored (Mouse Stat Jr, Kent Scientific) and all noted measurements were within the expected physiological range. For awake imaging, animals were trained to be head fixed for at least two weeks before the imaging session using sweetened condensed milk as treat.

### 4.8. *in vivo* experiment and data processing

#### 4.8.1. Experimental setup

Agarose phantom (no scattering) was used to fill the cranial window, which serves as the acoustic matching medium between a water container and the mouse brain. The bottom of the water container was covered with a thin clear film preventing water leakage. To maintain the brain temperature of experimental animal, degassed warm water (37° ± 1°) was circulating through the water container and, along with the agarose phantom, worked as the acoustic transmitting medium between the ultrasound transducer and the mouse brain, as shown in **Figure 4a**. An anteroposterior linear translating stage was used to carry the ultrasound probe to acquire data at different coronal planes.

For anesthetized imaging, the experimental animal was anesthetized by isoflurane through a nose cone while the body temperature was maintained at 37° with a homeothermic blanket control unit (Harvard Apparatus) and its head was fixed by a stereotaxic frame. For awake imaging, the experimental animal head was fixed by attaching the head-bar to a customized mount and the animal was treated with milk every ~30 min.

#### 4.8.2. In vivo validation

For in vivo validation, animals were anesthetized with isoflurane and the body temperature was maintained at 37°. vUS data was first acquired at different coronal planes and followed by microbubble injection for ULM/vULM imaging for each coronal plane. 0.03 ml commercial microbubble suspension (5.0-8.0× 10^8^ microbubbles per ml, Optison, GE Healthcare, Milwaukee, WI) was administered through retro-orbital injection of the mouse eye. The vULM map was rescaled to have the same pixel size (25×25 *μm*^2^) as vUS map. For a fair comparison, both the vULM and the vUS measurements were applied with a spatial mask that ensures nonzero valued pixels for both vUS and vULM measurements.

#### 4.8.3. Whisker stimulation

N=3 mice were trained and used for the whisker stimulation experiment. An air puffer machine (Picospritzer III, Parker Inc.) was used for the whisker stimulation experiments. The outlet of the air tube was placed ~15 mm behind the whiskers. Two stimulation patterns were used in this study: the first stimulation pattern (**Figure 4** and **Figure S5a**) consisted of 30 s baseline and followed by 10 trials of 15 s stimulation and with a 45 s interstimulus interval, and the second stimulation pattern (**Figure S5b**) consisted of 20 s baseline and followed by 10 trials of 5 s stimulation and with a 25 s interstimulus interval. A motion correction method was used to replace the signal value at strong motion time points with the median value of adjacent time points. The stimulation frequency was 3 Hz.

The whisker stimulation activation maps were calculated as the correlation coefficient *r* between the blood flow velocity *v*(*z*, *x*, t) and the temporal stimulus pattern *S(t)*.

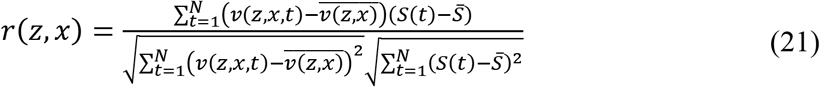

where,

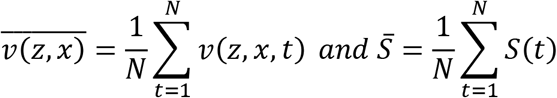

where, *N* is the total acquisition. The correlation coefficient was transformed to *z* score according to Fisher’s transform (**Equation 16**) and the level of significance was chosen to be *z>4.43* (*p* < 0.001, one tailed test), which corresponds to *r* > 0.2.

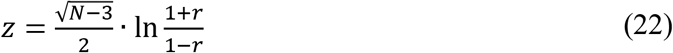

## Supporting Information

Supporting Information is available from the Wiley Online Library or from the author.

## Acknowledgements

The authors acknowledge funding from NIH R01-EB021018, R01 NS108472, and R01-MH111359.

## Authors contributions

J.T. and D.A.B. conceived of the technology and designed this study. J.T., D.D.P., T.L.S. and D.A.B. developed the theoretical model and analyzed the results. J.T. derived the theoretical formula, developed the data processing method, constructed the experimental setup, carried out experiments, and wrote the manuscript. K.K., E.E. and B.L. developed the surgical protocol for chronic imaging on awake mice, carried out animal experiments and analyzed the results. J.T.G designed the head bar. D.A.B. supervised this study. All authors discussed the results and contributed to the final version of the manuscript.

## Supporting Information

### I. Supplementary Figures

**Figure S1.**
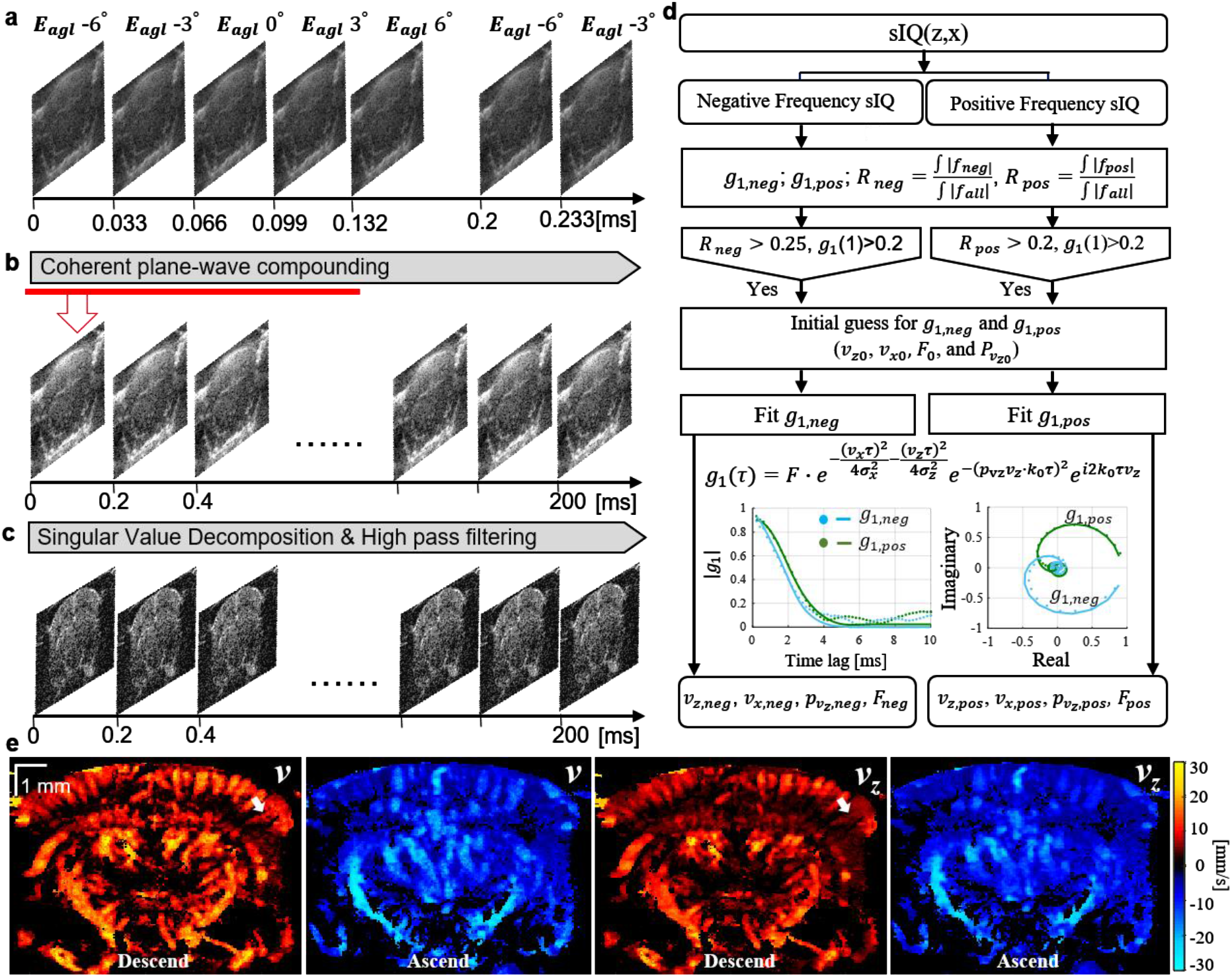
vUS implementation and data processing. (a) Ultrasound pulse & acquisition sequence. (b) Coherent plane-wave compounding were performed on the 5 tilted emission angle frames and produced a compounded image at a frame rate of 5 kHz. (c) Clutter rejection were performed to remove static background and bulk motion signal components. (d) Negative and positive frequency components of a measurement voxel are processed separately for *in vivo* data vUS processing; dots: experimental data; solid lines: fitting results. (e) Descending and ascending blood flow velocity maps reconstructed by vUS of a coronal plane (~Bregma −2.18 mm) of a mouse brain.

**Figure S2.**
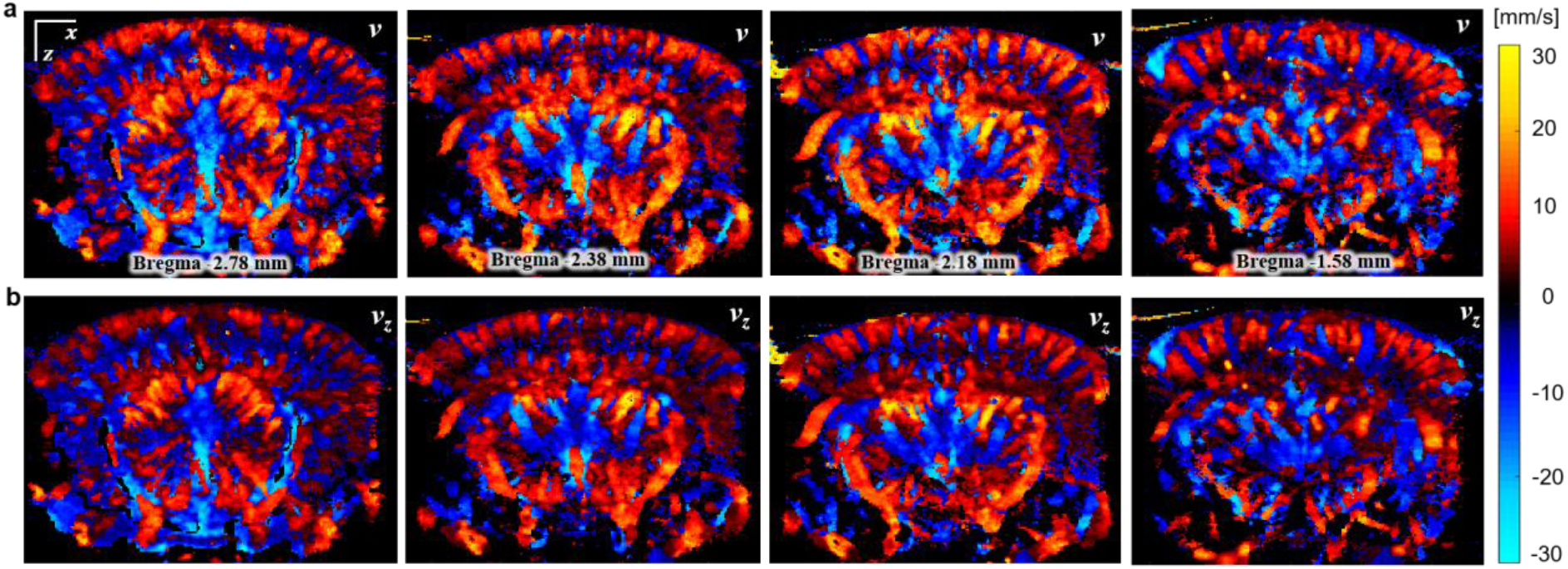
Total velocity (a) and axial velocity (b) obtained with vUS at different coronal planes of a mouse brain. Descending flow velocity map was overlapped on ascending flow velocity map.

**Figure S3.**
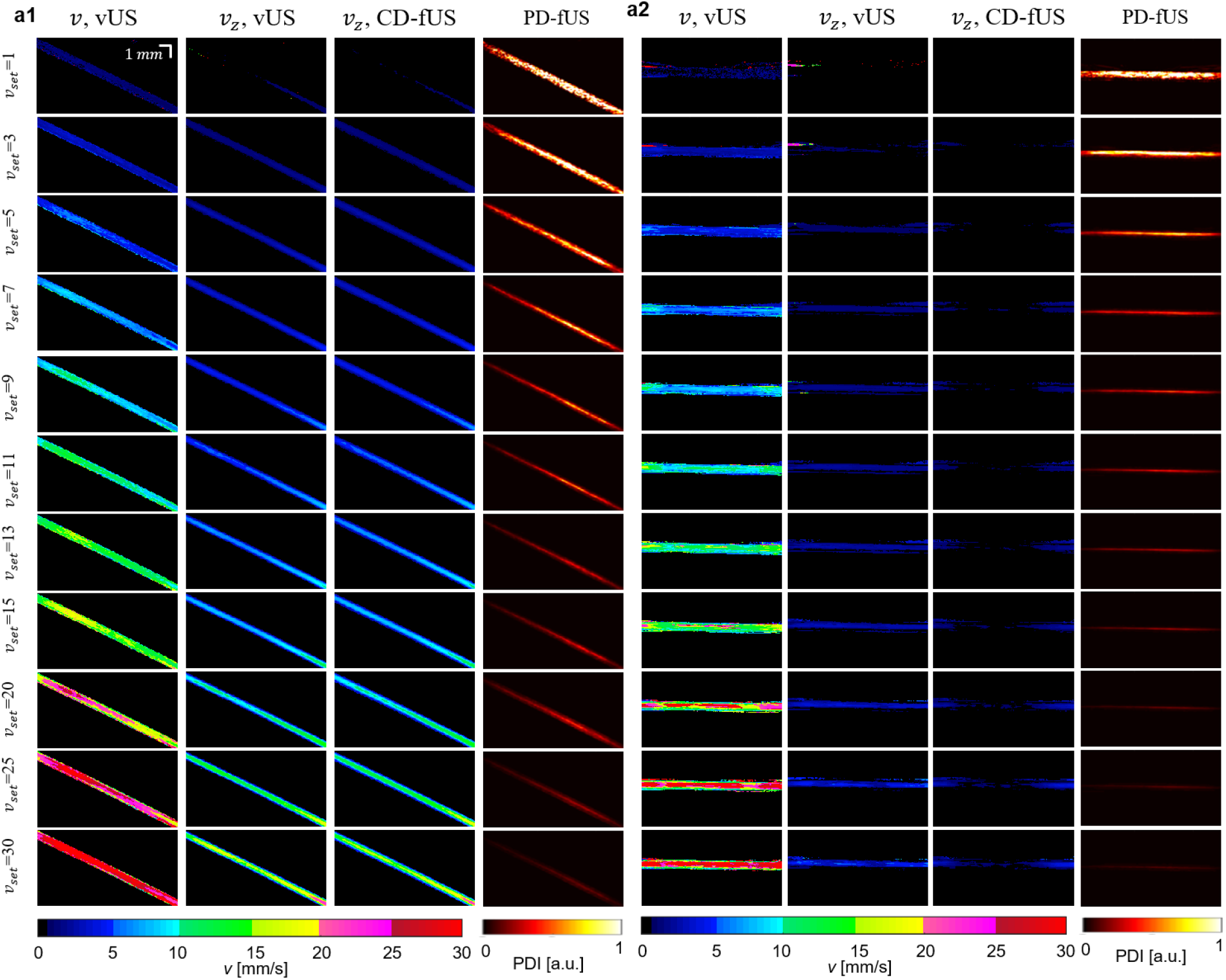
Phantom experiment validation and comparison. (a) Results for angled flow phantom experiments. (b) Results for transverse flow phantom experiments. vUS is able to accurately measure both axial and transverse velocity components while CD-fUS is not capable of measuring the transverse flow velocity component. In addition, vUS is able to accurately differentiate the axial velocity component from the transverse velocity component given its ability to determine flow direction. Compared to PD-fUS, vUS measured velocity has a linear relationship with the preset speeds, while the PD-fUS measured signal decreases nonlinearly with increasing preset speed.

**Figure S4.**
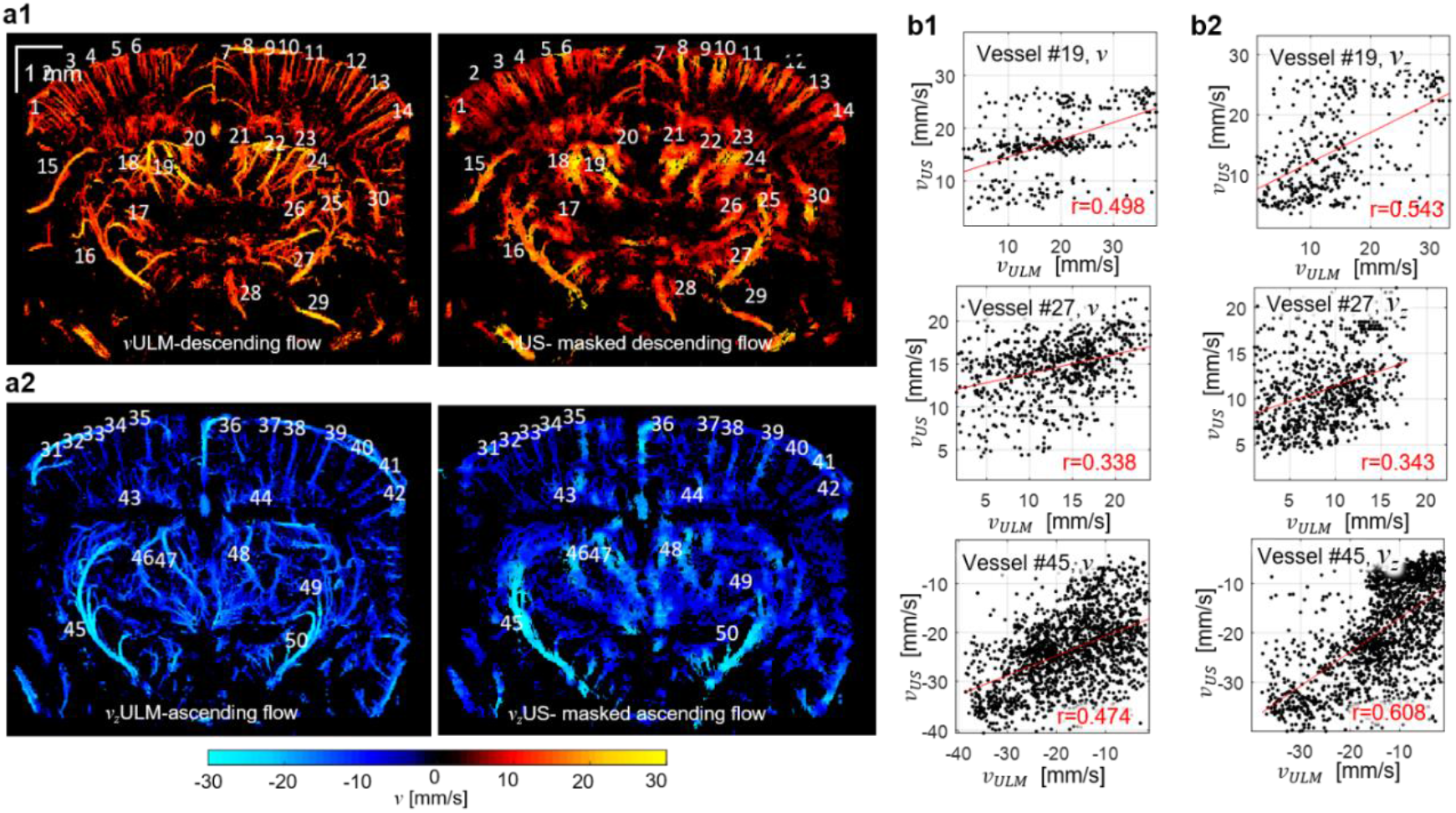
*in vivo* validation by comparing vUS with vULM. (a) The numbers show the indices of selected vessel for vessel-to-vessel comparison between vUS and vULM. (b1) Scatter plots of total velocity of three representative vessels show the pixel-to-pixel correlation between vULM and vUS. (b2) Scatter plots of axial velocity of three representative vessels show the pixel-to-pixel correlation between vULM and vUS.

**Figure S5.**
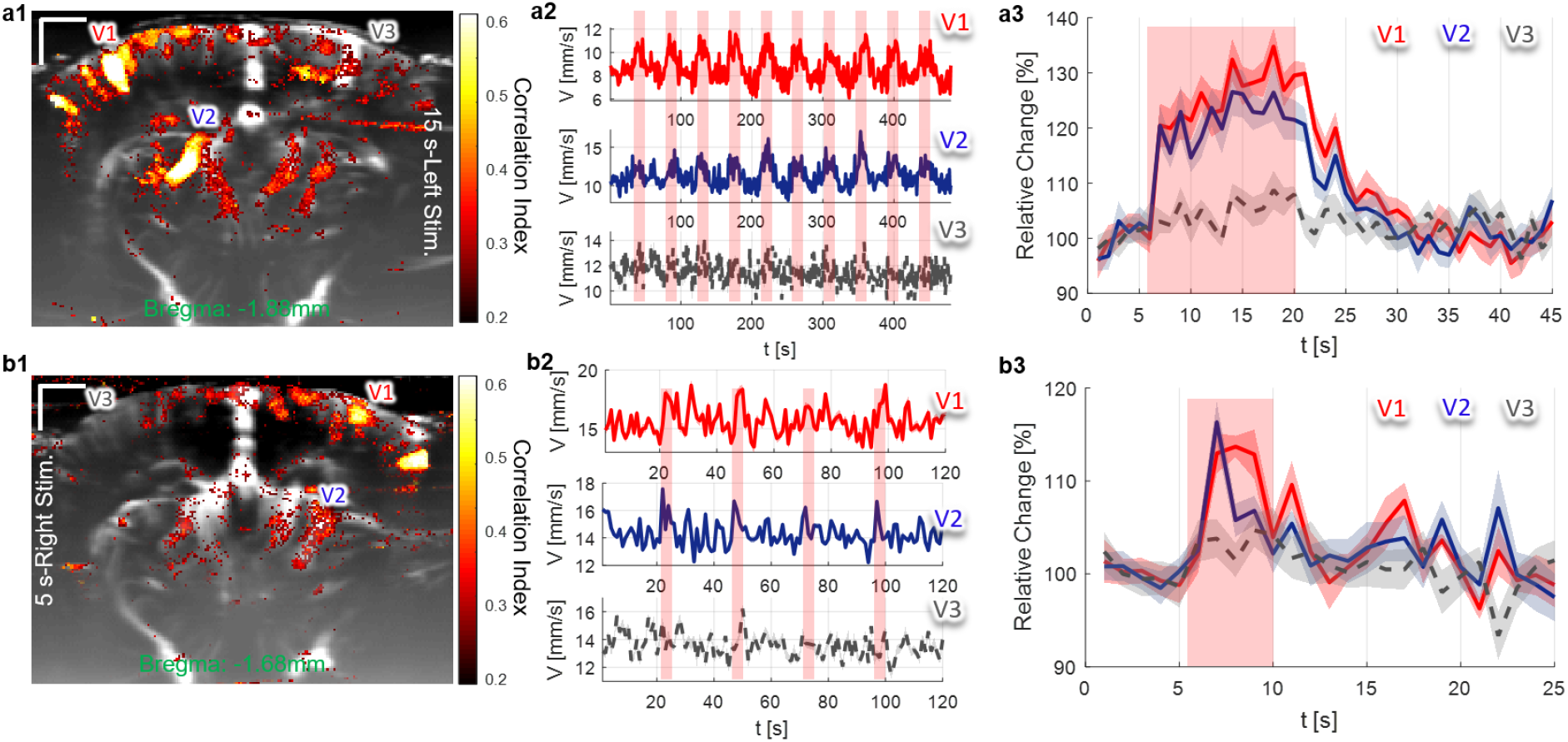
Representative whisker stimulation results. (a) Results of 15 seconds left side whisker stimulation; (a1) Activation map; (a2) Blood flow velocity time courses for the three vessels marked in (a1); (a3) 10 trials averaged relative response of the three vessels. (b) Results of 5 seconds right side whisker stimulation at Bregma ~ −1.58 mm; (b1) Activation map; (b2) Blood flow velocity time courses for the three vessels marked in (b1); (b3) 10 trials averaged relative response of the three vessels.

### II. Function description for vUS data processing

**Note**: vUS data processing code and example data is available from: <u>**Supplementary Code**</u>.

#### A. vUS data processing for *in vivo* data

##### A.1. main function

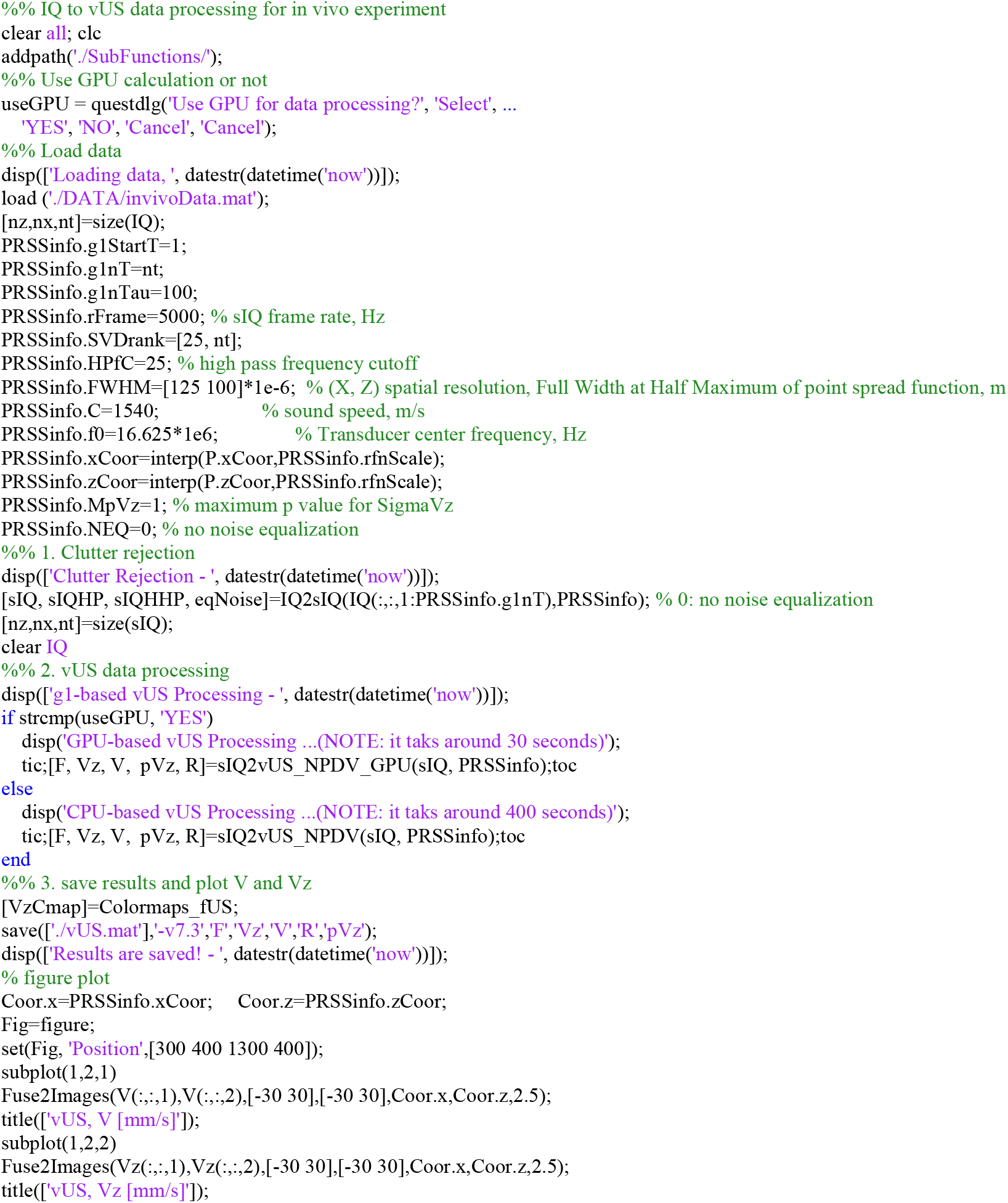

##### A.2. function IQ2sIQ

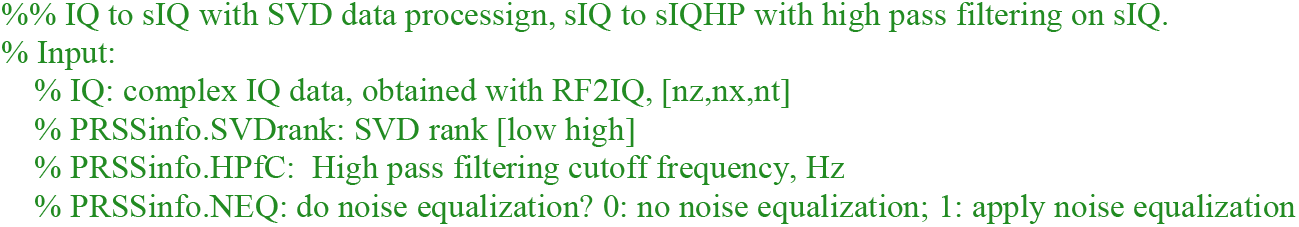
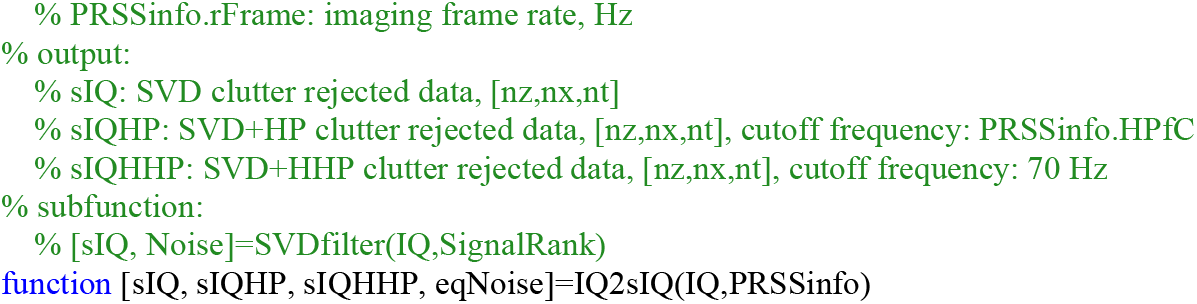

##### A.3. function sIQ2vUS_NP_DV

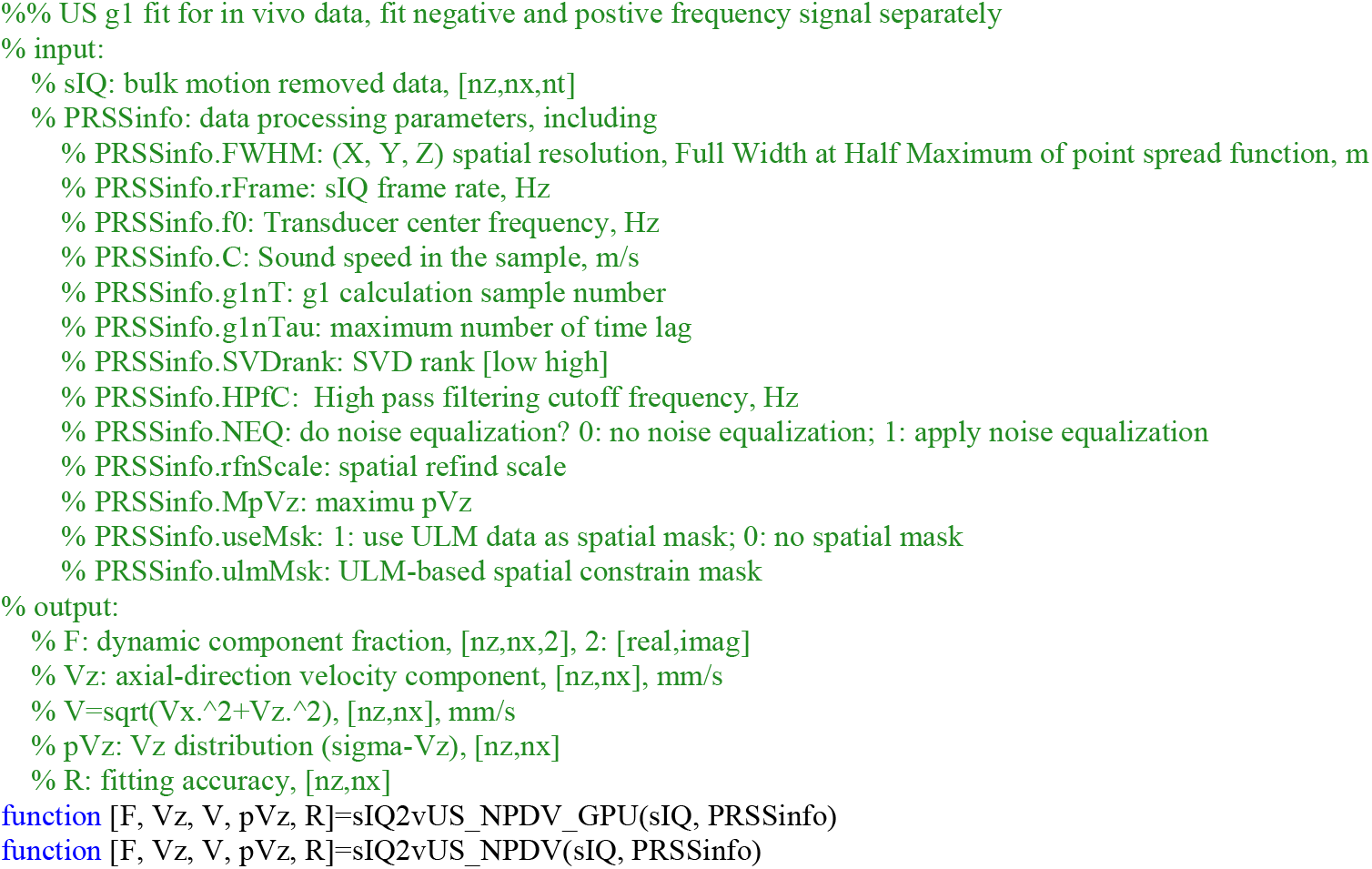

### B. vUS data processing (SV model) for phantom data

#### B.1. Main function

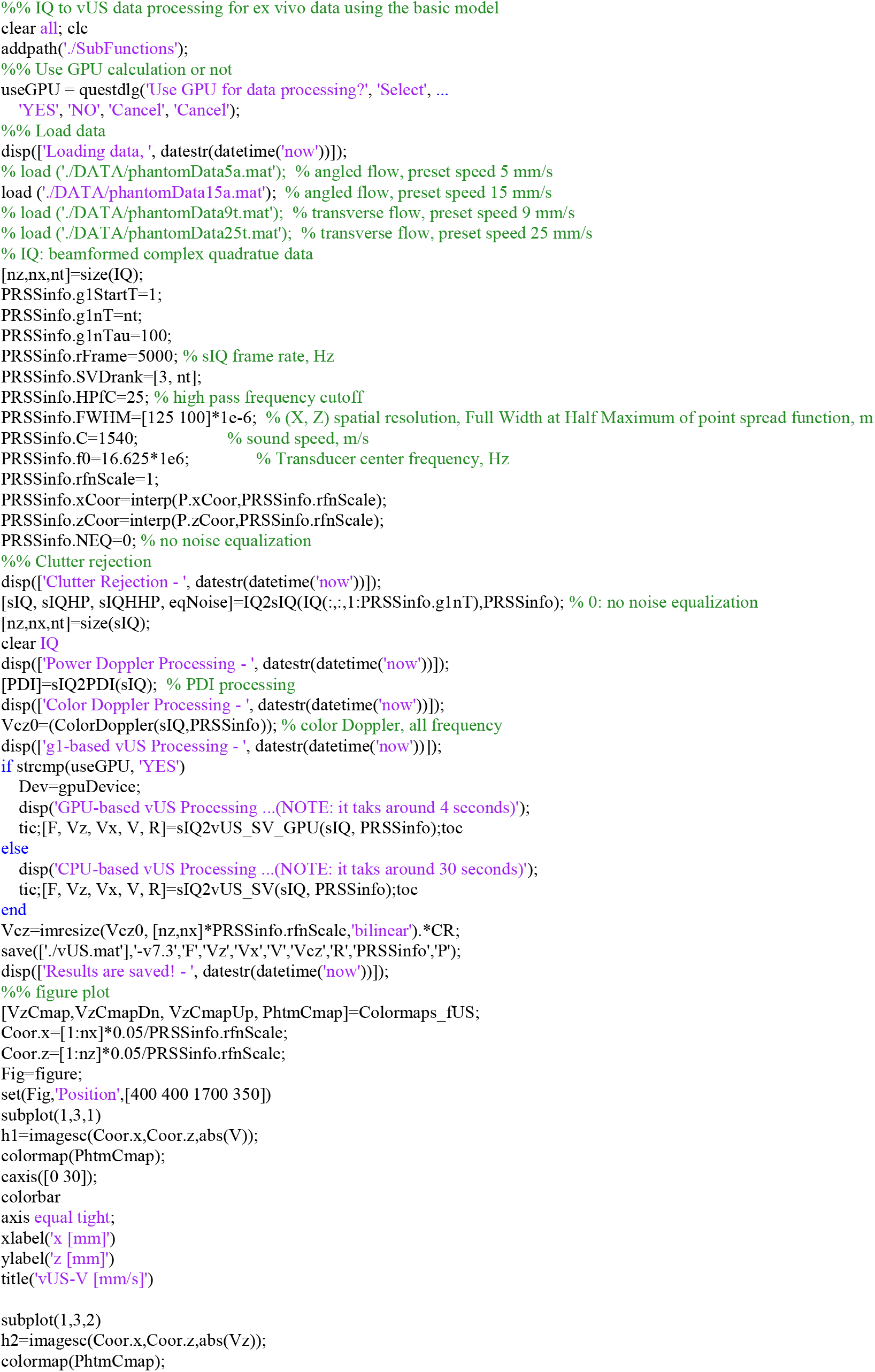
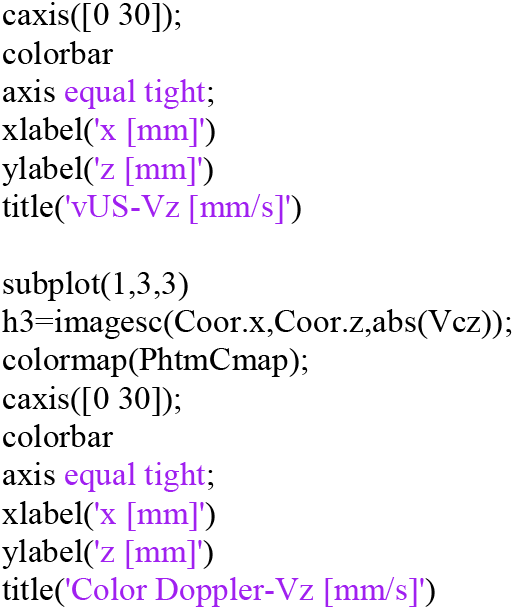

#### B.2. function sIQ2vUS_SV

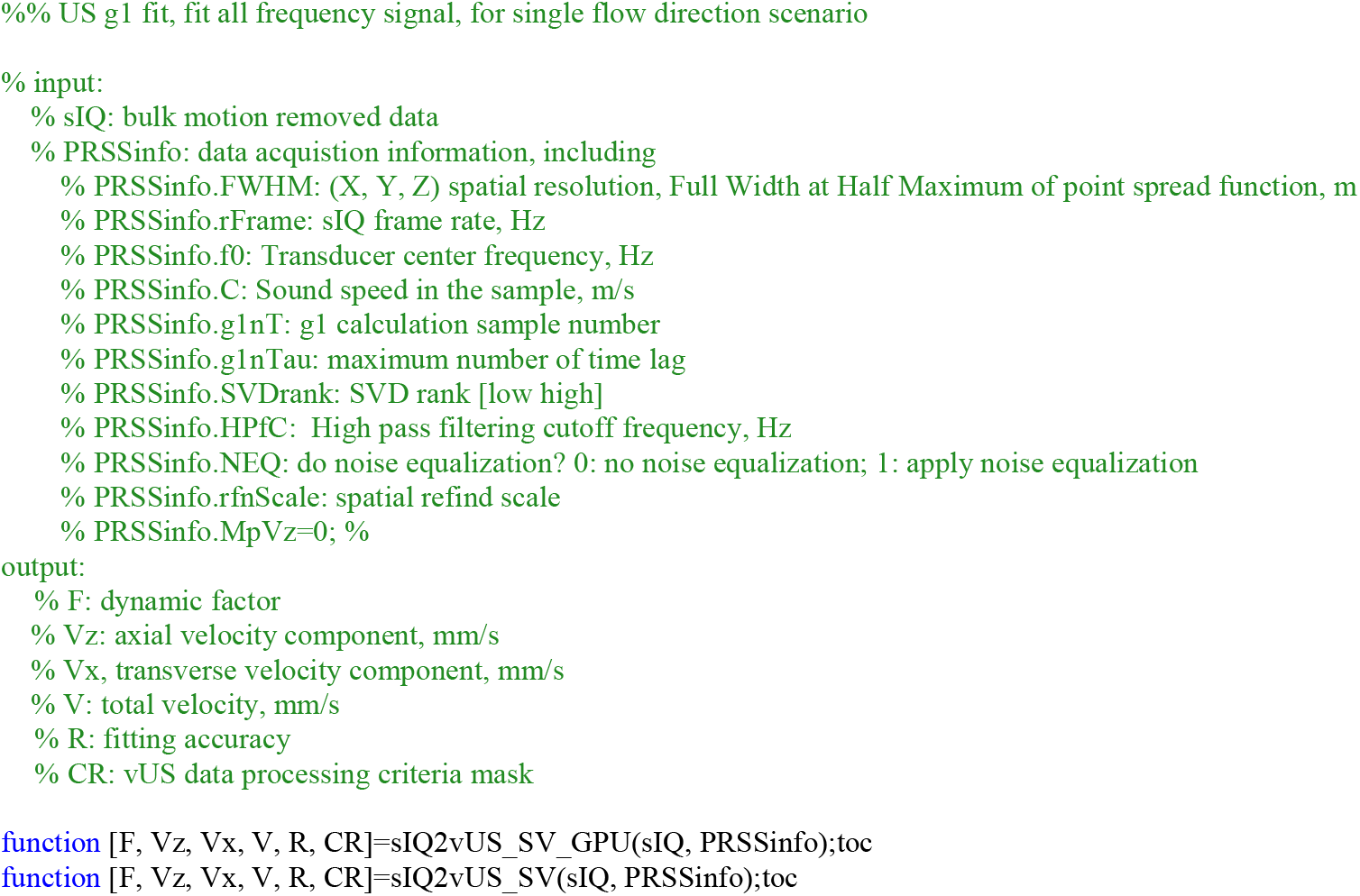

#### B.3. function ColorDoppler

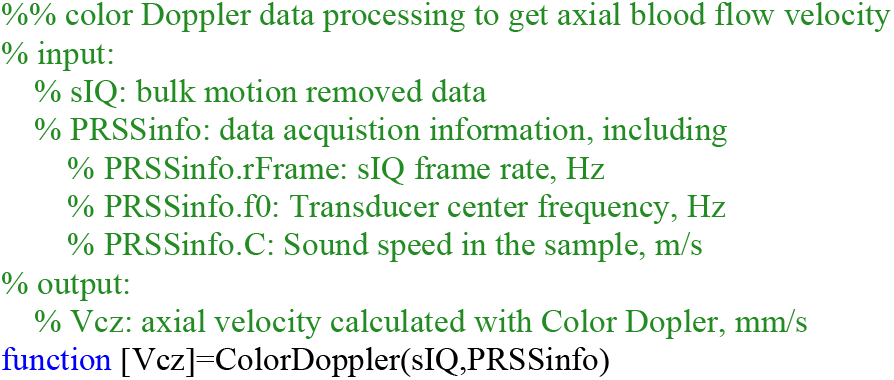

